# Multiscale intraspecific variation and coordination of hydraulic traits in silver fir

**DOI:** 10.64898/2026.05.28.728426

**Authors:** Belén Acuña-Míguez, Alice Copie, François Lefèvre, Maurizio Mencuccini, Ivan Scotti, Hervé Cochard, Sylvain Delzon, Arsène Druel, Bruno Fady, Florence Jean, Caroline Scotti-Saintagne, José M. Torres-Ruiz, Nicolas K. Martin-StPaul

## Abstract

- Intraspecific variability in hydraulic and morphological traits may alter projections of forest vulnerability to climate change. Recognizing multiscale nature is essential, as trait relationships within populations may not reflect those across the species’ range.
- We assessed variability in 11 traits in 10 natural populations of *Abies alba* across environmental and phylogeographic gradients to evaluate potential impacts on drought vulnerability. We quantified trait variation within and among populations, tested effects of climate, phylogeny and local factors, and evaluated trait coordination across scales.
- Hydraulic safety traits and wood density showed low variability, indicating strong constraints. In contrast, water-use and efficiency traits were highly variable. Aridity influenced several traits, but reduced variance was detected only for leaf residual conductance and succulence consisted with stabilising selection. Trait coordination was weak within populations than among populations.
- Overall, *A. alba* combines constrained hydraulic safety traits with variable water use traits. This may buffer drought impacts but limits shifts in hydraulic safety margins, potentially increasing risks of hydraulic failure, especially in humid environments where populations lack the capacity to tolerate prolonged dry periods. Thus, our findings highlight the need to account for scale-dependent trait variability, coordination, and their underlying drivers when predicting species’ adaptive capacity to drought.

## Introduction

Water transport, a key feature for plant productivity and ecosystem functioning, is projected to be altered by warming temperatures, declining water availability and intensifying drought regimes (Mencuccini et al., 2019). In woody plants, there is a broad variation in resistance to cavitation across biomes and among species within biomes, and part of this variation is explained by phylogenetic legacy (Maherali et al, 2004). Many tree species operate close to critical hydraulic thresholds at which xylem cavitation causes rapid losses of conductivity (Choat et al., 2012). Under intensive drought, proximity to these cavitation thresholds predisposes trees to hydraulic failure, ultimately resulting in the widespread forest decline and mortality reported across regions (Allen et al., 2010). Hydraulic and morphological traits govern transpiration and water transport and have emerged as primary predictors of drought-related mortality across ecosystems (Anderegg et al., 2016; Powers et al., 2020). As drought progresses, plants experience a sequence of water-stress thresholds that constrain key physiological processes, each tightly associated with specific hydraulic traits (Choat et al., 2018). Substantial variability in these hydraulic traits has been documented across species and environments. While many studies have focused on interspecific variability in hydraulic traits, an increasing body of evidence highlights the importance of intraspecific variation (Anderegg, 2015; Ramirez-Valiente, Poyatos, et al., 2025). Intraspecific variability in hydraulic traits across populations and environmental gradients can approach, and sometimes even exceed, the magnitude of variability observed at the whole community level (Garcia et al., 2022). Critically, such intraspecific variability can be large enough to influence drought survival and mortality risk and alter projections of forest vulnerability under climate change (Anderegg et al., 2019; López et al., 2021). As a result, evaluating species vulnerability to drought requires accounting for this variation, as it can significantly influence estimates of drought risk under changing environmental conditions.

The extent of intraspecific variation depends on the scale at which it is examined. At the within-population scale, phenotypic differences reflect microenvironmental heterogeneity, ontogeny, competition and within-population genetic diversity (Albert et al., 2010; Benavides et al., 2021). Fririon et al. (2023) found that the within-population phenotypic variance of drought sensitivity and the within-population phenotypic correlation between vigour and drought sensitivity were both partly explained by the local environment. At the among-population scale, variation reflects both broad environmental contrasts across the species’ range and the evolutionary history of populations (Ohsawa & Ide, 2008). Thus, these multiscale sources of variation can shape trait variation differently across scales. The magnitude of intraspecific variation is also strongly trait-and species-dependent (Vilà-Cabrera et al., 2015; Rosas et al., 2019). Some traits, such as xylem vulnerability to cavitation, exhibit relatively low variability and appear weakly plastic and evolutionarily constrained at the species level (Lamy et al. 2011, 2014; Ramirez-Valiente, Poyatos, et al., 2025), whereas others, such as hydraulic conductivity, Huber value and residual minimum conductance show higher variability (Rosas et al., 2019; Wang et al., 2025). Recognizing the multiscale nature of intraspecific variation and the differences in magnitude is essential for interpreting the trait-specific patterns of hydraulic diversity observed across species.

Intraspecific variability in functional traits can arise from genetic diversity within and among populations, environmental variation, phenotypic plasticity—the ability of a genotype to adjust trait expression in response to environmental signals—or genotype x environment interactions (Alberto et al., 2013; Chevin et al., 2013; Blackman et al., 2017; Ramirez-Valiente, Sanchez-Martinez, et al., 2025). Common gardens or reciprocal transplant experiments are required to disentangle genetic and environmental effects on trait variation (Ramirez-Valiente et al., 2020; Fady & Rihm, 2022). In natural populations (i.e. trees growing in their native environments), the term “phenotypic adjustment” has been proposed (Ramirez-Valiente, Poyatos, et al., 2025) to describe variation in mean trait values across environments that reflects local environmental conditions. Moreover, considering not only the variation of trait means but also the variation of within-population variances and correlations can provide insights into environmental and evolutionary factors shaping trait variability across scales (Ahrens et al., 2021; Anderegg et al., 2021). Notably, traits evolve together, and increasing evidence shows that drought resistance emerges from coordinated suites of traits rather than from single traits alone (Choat et al., 2018). Trait coordination can vary across environments (Benavides et al., 2021) and may reflect underlying genotypic differentiation (Sanchez-Martinez et al., 2024). Because the processes shaping phenotypic variation differ across spatial scales, the structure of trait covariation may also shift between within- and among-population levels (Alonso & Herrera, 2001). Consequently, the relationships among traits observed within populations may not necessarily reflect the patterns that emerge when comparing populations across the species’ range.

Different strategies have been proposed to disentangle genetic, environmental, and/or genotype–environment interaction effects on traits in natural populations among species. Halliwell et al. (2025) reviewed the use of phylogenetic mixed models (PMMs) as an approach to partitioning phylogenetic and non-phylogenetic components of single and multivariate trait variation. At the intraspecific level, phylogenetic information can be simplified to a relevant clustering level and included as a qualitative grouping factor, thereby simplifying the PMM approach. To cope with confounding effects between phylogeny and environment, Sánchez-Martínez et al. (2024) proposed a Bayesian PMM framework. However, in cases where environmental variation is not correlated with phylogeny, the separation of these effects is further facilitated. Thus, complementary analytical approaches that integrate phylogenetic structure, environmental conditions, and additional sources of variation, together with analyses of variance across populations in contrasting environments and trait covariation within and among populations of the same species, may provide a more general and robust framework for studying intraspecific variability in functional traits.

*Abies alba* Mill. (the silver fir) is a temperate forest tree species adapted to cold environments, yet its potential to persist under future climate change remains unclear, with contrasting responses reported across populations (Büntgen et al., 2014; Gazol et al., 2015; Oggioni et al., 2022). Moreover, some populations of this species acted as climate refugia during glacial periods (Liepelt et., 2009; Scotti-Saintagne et al., 2021, Pèlachs et al., 2026), providing long time windows for local adaptation to occur (Liepelt et al., 2009). These two factors make *A. alba* a suitable model for studying intraspecific variability in hydraulic traits across its natural range. Here, we assess the adaptive potential for drought response in *A. alba* by analysing genetic structure and phenotypic variation in hydraulic traits and their coordination across 10 populations spanning environmental and phylogeographic gradients across its natural range. Specifically, we aim to: (i) to quantify the magnitude of phenotypic trait variability in *A. alba* in order to identify which traits are more evolutionary constrained and which are more variable; (ii) to quantify the role of phylogenetic legacy and local environment on the patterns of trait means (phenotypic adjustment) and within-population variances; (iii) to characterise phenotypic trait coordination within and among *A. alba* populations, and (iv) to examine whether population-level differences in coordination patterns are structured by phylogenetic clusters, environmental conditions or their interactions.

## Material and methods

### Study plots

Nineteen natural forest plots, belonging to the pan-European EUFORGEN Genetic Units Conservation (GCU) network (www.euforgen.org, eufgis.org), were selected across *A. alba*’s natural range (Fig. 1). GCUs were selected based on bioclimatic representativeness with an over-representation of marginal areas.

**Figure 1.**
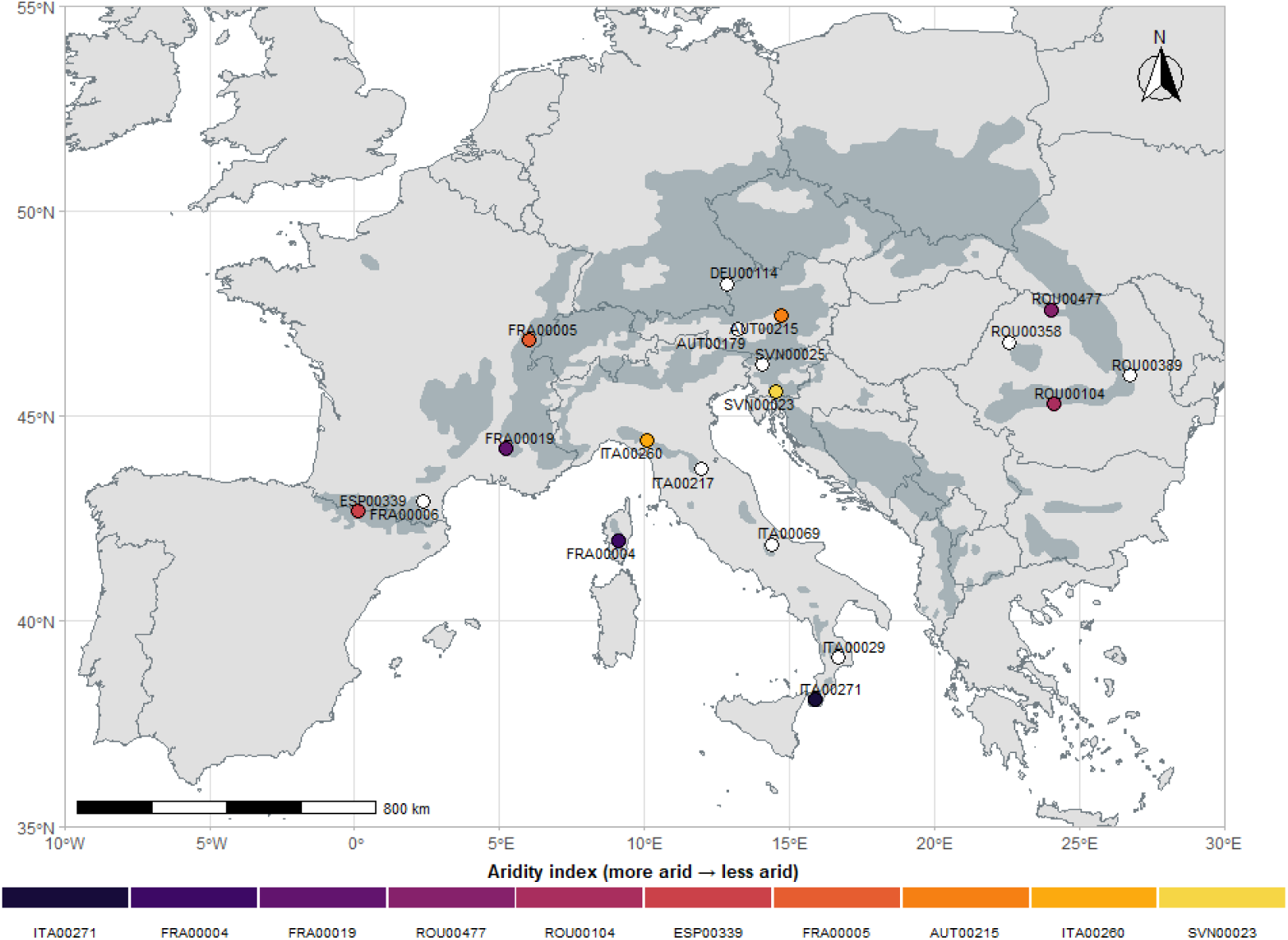
Geographic distribution of *A. alba* populations across Europe. Coloured symbols represent the aridity index for the 10 populations selected for trait measurements, whereas non-coloured symbols correspond to additional populations included for genetic analyses. The natural distribution range of *A. alba* is shown in dark grey.

### Genetic data

We took the genetic variation data, consisting of 66940 Single Nucleotide Polymorphism (SNP) loci, characterised in 25 individuals sampled within a single, environmentally homogenous patch in each GCU, and recorded in a VCF (Variant Calling Format) file (Pinosio et al., 2025). Selected trees were dominant or codominant adults, belonging to the same age class, separated by a minimum distance of 30 meters (Mariotte et al., 2025 for further details).

### Sample trees for phenotypic analyses

Among the nineteen plots, ten populations spanning the environmental gradient of *A. alba* were selected for phenotypic analyses (Fig. 1). In each population, elevation and slope were also recorded, and stand density and stand basal area were measured including trees of all species in the plot (Table 1, supplementary material). We computed an aridity index (Zomer, et al., 2022) for the period 1970-2000 period at 30’’ spatial resolution— defined as Mean Annual Precipitation divided by Mean Annual Reference Evapotranspiration (lower aridity index values indicate higher aridity)—and the precipitation of the warmest (bioclimatic variable BIO18) from the WorldClim database (Fick & Hijmans, 2017) which has been shown to be an indicator of drought resistance in the genus *Abies* (Copie et al., 2025).Ten individuals per plot were selected, preferentially from the 25 individuals previously sampled for genetic analyses. We georeferenced each individual and measured biometric characteristics, age and basal area assessment of living and dead trees around each sample tree (Table 1).

**Table 1.** Mean and standard deviation (SD) of CBH (circumference of the trunk at 130 cm, cm), Ht and Hbc (Height and basal crown height, respectively, m), LgCrown (Length of the crown= Ht-Hbc), CrownArea (the footprint of the tree crown area estimated by 4 length projected in the 4 directions: North-East-South and West for each Functional Subset tree, m), Age (years) g_living and g_dead (basal area assessment for living and dead tree around each tree, respectively, according to the chosen band, m2/ha) of sampled trees. Different letters show significant differences among populations for each characteristic.

|  | ITA00271 | FRA00004 | FRA00019 | ROU00477 | ROU00104 | ESP00339 | FRA00005 | AUT00215 | ITA00260 | SVN00023 |
| --- | --- | --- | --- | --- | --- | --- | --- | --- | --- | --- |
| CBH | 157.7 ± 32.05 (ab) | 141.9 ± 25.75 (ab) | 149 ± 33.27 (ab) | 212.7 ± 33.39 (cd) | 235.1 ± 67.98 (c) | 127.3 ± 28.16 (b) | 225.3 ± 38.22 (cd) | 181.3 ± 28.99 (ad) | 120.6 ± 24.91 (b) | 225.4 ± 31.26 (cd) |
| Ht | 19.66 ± 2.65 (ab) | 23.74 ± 3.19 (a) | 18.42 ± 2.43 (b) | 33.6 ± 4.21 (c) | 30.72 ± 4.25 (c) | 22.06 ± 3.05 (ab) | 33.88 ± 3.36 (c) | 34.02 ± 1.9 (c) | 19.28 ± 3.18 (ab) | 34.26 ± 3.52 (c) |
| Hbc | 7.58 ± 3.18 (a) | 13.56 ± 3.5 (bc) | 10.74 ± 3.13 (acd) | 20.98 ± 3.22 (e) | 15.45 ± 2.3 (bdf) | 9.97 ± 3.22 (ac) | 16.03 ± 3.75 (bf) | 8.98 ± 4.52 (ac) | 9.32 ± 2.92 (ac) | 19.56 ± 3.15 (ef) |
| LgCrown | 12.08 ± 1.29 (abc) | 10.18 ± 2.39 (bc) | 7.68 ± 1.53 (b) | 12.62 ± 2.56 (acd) | 15.27 ± 3.7 (ae) | 12.09 ± 3.23 (abc) | 17.85 ± 4.15 (e) | 25.04 ± 4.49 (f) | 9.96 ± 2.96 (bd) | 14.7 ± 4.11 (ce) |
| CrownArea | 34.37 ± 11.69 (a) | 38.7 ± 13.87 (a) | 44.34 ± 18.14 (ab) | 30.11 ± 9.87 (a) | 33 ± 17.49 (a) | 36.32 ± 12.96 (a) | 36.48 ± 20.67 (a) | 39.66 ± 24.56 (a) | 30.61 ± 7.66 (a) | 68.9 ± 28.12 (b) |
| Age | 106 ± 66.69 (abc) | 90 ± 18.68 (bc) | 162.3 ± 28.98 (de) | 116.11 ± 30.19 (cdef) | 164.2 ± 72 (d) | 53.2 ± 13.34 (b) | 107.8 ± 23.21 (be) | 157.3 ± 28.29 (ade) | 81.6 ± 18.72 (bf) | 144.4 ± 25.98 (cde) |
| g_living_ha | 33.8 ± 7.3 (a) | 56.8 ± 12.97 (b) | 18 ± 5.25 (c) | 17.95 ± 2.02 (c) | 16.25 ± 6.35 (c) | 48.6 ± 20.96 (b) | 13 ± 3.61 (c) | 13.05 ± 2.52 (c) | 44.6 ± 9.65 (ab) | 34.3 ± 7.82 (a) |
| g_dead_ha | 2 ± 1.63 (ab) | 5.2 ± 4.92 (a) | 4.6 ± 5.34 (ac) | 1.5 ± 2.04 (ab) | 2.1 ± 1.47 (ab) | 1.2 ± 1.69 (ab) | 0.0500 ± 0.1581 (b) | 0.6000 ± 0.7746 (bc) | 3.2 ± 3.55 (ab) | 0.4000 ± 0.8433 (b) |

### Study traits

We collected 2 or 3 long, leafy, sun-exposed branches (about 1 m long and 3 cm in diameter at the base) on each of the 10 trees. We subsampled each branch to collect shoots and leaves for the assessments of 11 traits used in the present study (Table 2; Mariotte et al., 2025 for further details; mean and standard deviations of each trait in each population are shown in table 2 supplementary material).

**Table 2.** Study traits, their units and function attributable to them.

| Trait | Unit | Function |
| --- | --- | --- |
| **Specific leaf area (SLA)** | m<sup>2</sup>.g<sup>-1</sup> | Refers to carbon gain relative to water loss. |
| **Wood density (WD)** | g cm<sup>-3</sup> | Carbon investment in trees.<br>Denser wood offers enhanced mechanical support and greater resistance to drought. |
| **P50** | MPa | Xylem water potential inducing 50% loss of hydraulic conductance. It represents water stress threshold for survival in conifers. The more negative the values, the greater the vulnerability to cavitation. |
| **Slope of the vulnerability curve at P50 (Slope)** | MPa<sup>-1</sup> | How fast xylem embolism occurs |
| **Turgor loss point (TLP)** | MPa | Leaf water potential at which leaf wilting occurs, considered a proxy for stomatal closure. More negative values indicate greater turgor maintenance and later stomatal closure (Martin-StPaul et al., 2017). |
| **Leaf residual conductance (g<sub>res</sub>)** | mmol m<sup>-2</sup> s<sup>-1</sup> | Loss of water through the cuticle and incompletely closed stomata. Measured between 80 and 60% RWC. |
| **Succulence (Suc)** | g m<sup>-2</sup> | Ratio of water mass at full hydration to leaf area. A proxy of water storage capacity in leaves (Blackman et al., 2019) |
| **Xylem specific hydraulic conductivity (K<sub>s</sub>)** | m<sup>2</sup> MPa<sup>-1</sup> s<sup>-1</sup> | K<sub>s</sub> was calculated by dividing K<sub>max</sub> (maximum xylem hydraulic conductivity initially obtained when the pressure was close to zero in the cavitron) by the length of the sample and divided by the cross-sectional area (Larter et al. 2017). |
| **Huber value (HV)** | cm<sup>2</sup> cm<sup>-2</sup> | Sapwood to leaf area ratio. It can be considered also as a proxy of hydraulic efficiency (Jacobsen et al., 2007; Markesteijn et al., 2011). |
| **Stomatal Margin Retention Index (SMRI)** | MPa m<sup>2</sup> s mmol<sup>-1</sup> | Indicator of the time required to cross the stomatal safety margin. |
| **Time to Hydraulic Failure (THF)** | Days | An indicator of vulnerability which is an output of the mechanistic model SurEau (Cochard et al., 2021). |

Below, we summarise methods used for each trait (for further details, Mariotte et al., 2025):

#### Vulnerability curves

We quantified xylem vulnerability to cavitation using a centrifuge-based method (cavitron) following Cochard (2002) by exposing branch segments to progressively more negative pressure at the centre of each sample while measuring hydraulic conductance. All measurements were taken using a standard cavitron (27 cm rotor) at the PHENOBOIS platform (INRAE, University of Bordeaux; Pessac, France). We expressed the reduction in conductance as percentage loss relative to maximum values (Kmax) and used these values to construct vulnerability curves, which we fitted with a sigmoidal model to estimate the water potential causing 50% loss of conductivity (P50) and the slope of the curve at P50.

We calculated hydraulic conductance (kh) as the ratio between water flow and the pressure gradient across the sample following Wang et al. (2014), and we obtained specific hydraulic conductivity (K_s_) by normalizing kh by the sapwood cross-sectional area and sample length (see Burlett et al., 2022 for further details).

#### Leaf turgor loss point

We collected at least 10 leaves from two branches and rehydrated them for 12–24 h at 4 °C in darkness with the petiole submerged in distilled water to ensure full saturation. Turgor loss point (TLP) was estimated by extracting leaf sap after membrane rupture and measuring osmotic potential at saturation using a freezing point osmometer (Advanced Instruments 3320 Micro-Osmometer). TLP was then calculated following following Barlett et al. (2012):

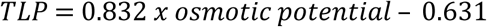

#### g_res_ and succulence

We computed leaf residual conductance from water losses measured under controlled conditions using the DroughtBox chamber method (Billon et al., 2020; Burlett et al., 2025). A subsample of three apical leafy shoots (at least 35-40 cm long) from each branch tree were selected. We rehydrated shoots and measured their weight and their basal diameter and length of the branch (used for estimated stem area). Afterwards, we installed them in a DroughtBox at 40% relative humidity and 30°C for monitoring changes in shoot weight every 5 minutes. We measured weight and water potential at the end of DroughtBox measurements. Finally, we separated the leaves and twigs and dried them in the oven for 48h at 60°C and measured their dry weight for twigs and for leaves. We estimated g_res_ following Burlett et al. (2025) by calculating the slope of mass loss per unit projected leaf area over time and dividing it by the vapor pressure gradient at a relative water content between 0.8 and 0.6. Leaf succulence was calculated as the difference between fresh mass and dry mass, divided by leaf projected area.

#### Huber value

We determined Huber value (HV) as the ratio between the leaf area subtended by a stem and the active xylem cross-sectional area of that stem (sapwood). We excluded dead or living bark, as well as the central pith or heartwood, and included only xylem rings that were active in water transport. To verify which rings were active, we applied a dye method (e.g., toluidine blue) to a subset of samples per species and used these observations to estimate the active sapwood area in the remaining samples.

#### Specific leaf area

We calculated specific leaf area (SLA) as the ratio between fresh projected leaf area and leaf dry mass. We randomly selected 30 to 60 leaves, scanned them, and measured their projected area using ImageJ software (Wayne Rasband, National Institutes of Health, Bethesda, MD, USA). We then oven-dried the samples for 48 hours at 60 °C and weighed them to determine leaf dry mass.

#### Wood density

We obtained wood density using a volumetric technique. We separated the bark from the xylem and determined the fresh volume of the wood by water immersion (Archimedes’ principle). We then oven-dried the samples at 70 °C for 48 h, recorded their dry mass, and calculated wood density as dry mass divided by fresh volume.

#### Stomatal Margin Retention Index (SMRI)

We calculated SMRI following Petek-Petrik et al. (2023) as:

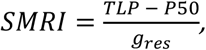

where TLP is used here as a proxy for water potential inducing stomatal closure.

#### Time to hydraulic failure

We used the SurEau model (version presented in Ruffault et al 2022) under constant climate conditions to estimate a standardized time to desiccation, a metric called Time to Hydraulic Failure (THF, e.g., Moreno et al 2024; Mas et al 2024). To do this, we followed the configuration procedure for SurEau used by Copie et al. (2025).

### Statistical analyses

#### Genetic distance matrix and population clustering

We computed allele frequencies in each GCU and for each SNP using vcftools (Danecek et al. 2011). We collected the frequency of the ALT (“alternate”) alleles from vcftools output.We computed the population-pairwise variance–covariance matrix of allele frequencies using the base R function *cov*(), which summarizes multivariate genetic affinity among populations. Because this covariance matrix represents similarity rather than distance, we converted it into a dissimilarity matrix using a monotonic transformation that preserves the rank order of affinities while expressing them on a dissimilarity scale appropriate for hierarchical clustering. The resulting dendrogram is interpreted as an exploratory visualization of broad genetic relationships rather than as an evolutionary distance tree. Hierarchical clustering was performed using the hclust() function from *stats* package and the dendrogram was used to visualize major population groupings. These clustering-derived groups were subsequently used as categorical factors in the analyses.

#### Magnitude of trait variability

We quantified intraspecific variation (coefficient of variation, CV) at two levels. Within-population variability was estimated as the CV of individual trait values within each population (standard deviation divided by the mean × 100). Among-population variability was quantified as the CV of population mean trait values, calculated as the standard deviation of population means relative to their overall mean.

#### Phenotypic adjustment and environmental effect on within-population trait variation

Before analysing trait variation, we tested whether phylogenetic groups differed in their climatic context. We evaluated group differences using a MANOVA with precipitation of the warmest quarter and aridity index as joint responses, followed by univariate ANOVAs for each variable. These analyses showed no significant association between phylogeographic groups and climate. For this reason, we analysed the source of variation in each functional trait using Bayesian linear mixed-effects models fitted with the *MCMCglmm* package (Hadfield, 2010), which implements Markov Chain Monte Carlo (MCMC) sampling for hierarchical mixed models. Each model included as random effects the phylogenetic groups and the populations nested in their phylogenetic groups. All models used Gaussian errors and weakly informative priors. Fixed effects included potential drivers of trait variation: tree age, aridity index, precipitation of warmest quarter and a proxy of tree competition estimated by basal area of living trees around each individual, all scaled (z-scored) before modelling. The inclusion of either the aridity index or precipitation of the warmest quarter as a fixed effect was based on preliminary analyses of rank correlations between population mean trait values and climatic variables (Supplementary Material, Fig. 1), as well as on comparisons of the variance explained by fixed effects across alternative models including either the aridity index or precipitation of the warmest quarter (Supplementary Material, table 3; Fig. 2). Formally, the models can be expressed as:

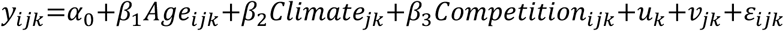

where 𝑦_𝑖𝑗𝑘_ is the trait value of individual i from population j belonging to phylogenetic group k; 𝛼_0_ is the intercept; 𝛽_1_- 𝛽_3_ are fixed-effect coefficients for tree age (Age), climatic variable (Climate) and tree competition (Competition) respectively; 𝑢_𝑘_ represents the random effect of phylogenetic group; 𝑣_𝑗𝑘_ is the random effect of population nested within phylogenetic group; and 𝜀_𝑖𝑗𝑘_ is the residual error.

**Figure 2.**
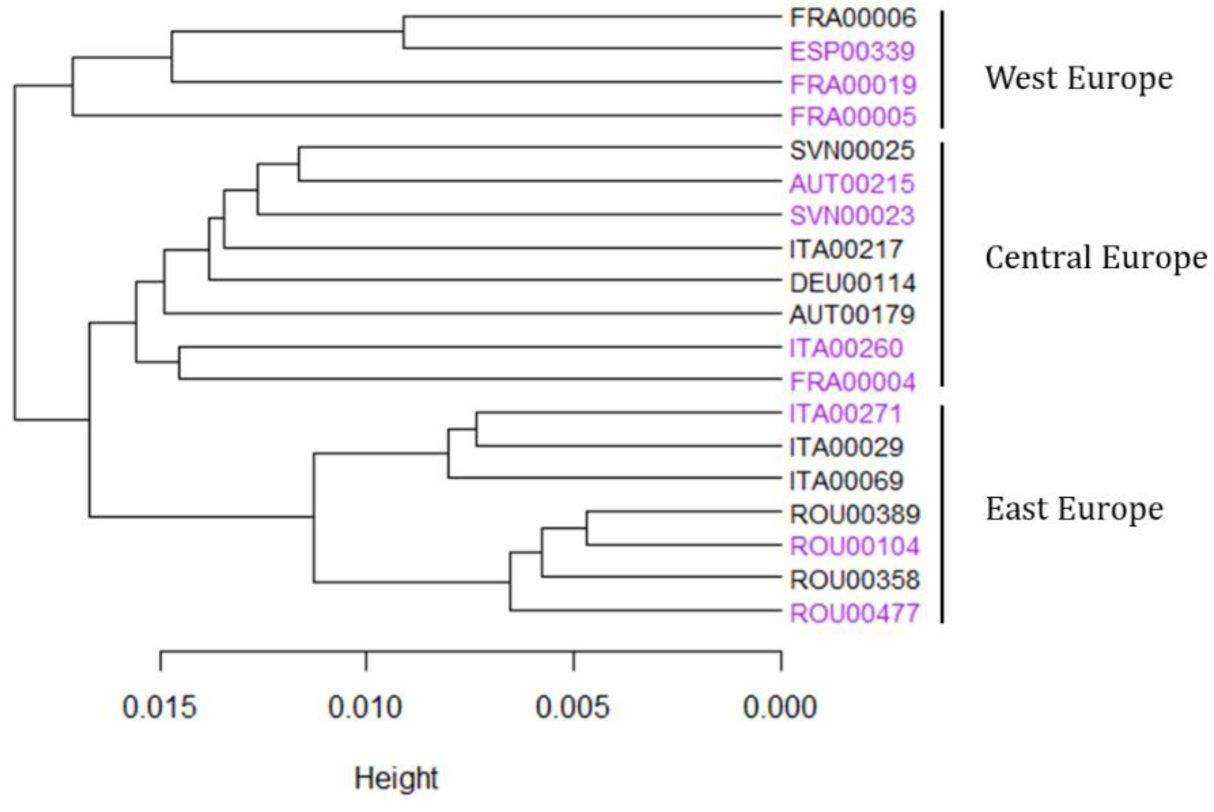
Hierarchical clustering of *A. alba* populations based on allele-frequency covariance. Coloured labels indicate the ten populations selected for trait measurements. Branch height represents the dissimilarity among populations.

Models were run for 50000 MCMC iterations, with a burn-in period of 10,000 iterations and a thinning interval of 10, resulting in a posterior sample of 4000 posterior samples per model. Convergence was verified visually by inspecting trace plots of both fixed and random effects parameters, which showed stable mixing and no apparent trends. Each trait was analysed independently with a Gaussian error distribution and g_res_, WD, SLA and SMRI were log-transformed before analysis to meet normality.

We extracted the posterior mean slope, along with 89% credible intervals to evaluate the effect size and direction following (Barry et al., 2025). We then used the posterior probability of direction (PD), defined as the proportion of posterior samples on the same side of zero, as an evidence metric for directional effects (Makowski et al., 2019) and categorized evidence strength as moderate (PD > 0.9), strong (PD > 0.95), or very strong (PD > 0.975). To describe how variance was distributed across hierarchical levels of the models, total phenotypic variance was decomposed into contributions from fixed effects, phylogenetic groups, populations nested within phylogenetic groups, and residual variance. For each posterior sample, the variance associated with each component was expressed as a proportion of the total model variance. Reported values correspond to posterior means, with uncertainty summarized using 0.89 credible intervals.

To better understand the sensitivity of plant traits in the SurEau model, variance-based sensitivity analyses were performed using the Sobol method (Sobol, 2001). This approach assesses the impact of parameter variation on model outputs and provides total-order indices that quantify each parameter’s contribution to output variance. Parameter ranges were defined based on the minimum and maximum values measured at each site (or across all sites for pooled *A. alba*). Analyses were conducted using the *sensobol* R package (version 1.1.5), computing second-order indices with an initial sample size of 10,000 values per parameter, resulting in approximately 230,000 simulations per site. (Supplementary material, Fig. 3)

To test for an environmental control on within-population phenotypic variances, we restricted the analysis to the six traits for which the relationship between population means and SD was not significant, indicating that among-population differences in within-population variability were not simply a mathematical consequence of changes in the mean (Supplementary Material, Fig. 3). We then analysed, for each trait, the relationships between population trait SD and CV (z-scored values) and the aridity index as potential environmental drivers of the intensity of selection, using Spearman’s rank correlations to capture any type of monotonous relationship (stats R package; R core team (2025)). We also analysed the correlations of trait variation and precipitation of the warmest quarter (Supplementary Material, Fig. 4). This analytical framework served as a first attempt to identify signals of stabilising selection on the traits (stabilising selection in broad sense, i.e. reducing the variance with or without changing the mean). If within-population variation differs among population beyond what is expected from mean-variance scaling and this variation is correlated with aridity index, we suspect a possible signal of selection. Here, we used two measures of within-population trait variation, SD and CV, considering that selection typically affects within-population SD while phenotypic adjustment can also change the mean, resulting in different patterns for both measures.

#### Trait coordination among and within populations

Finally, we tested trait coordination among and within populations. First, we z-standardized all traits prior to analysis. Pairwise correlations were decomposed into within-population and among-population components using *statsBy()* function from the *psych* R package (Revelle, 2025). The within-population correlation matrix estimates the association among traits at the individual level, after controlling for differences in population means. This is achieved by removing the between-population variation (i.e., subtracting each population mean for each individual trait value) and then calculating correlations across all individual residuals pooled across populations. The among-population matrix estimates correlations across population means (i.e., covariation between populations averages).

Considering both matrices (within and among population), we calculated three quantitative measures of trait covariation degree following Benavides et al. (2021): the edge density (ED), assessed as the ratio between the number of significant correlations (edges) and all possible pairwise trait combinations; the functional variability shape (FS) as the variance of the eigenvalues of the trait correlation matrix which considered a phenotypic integration index and modularity of the network (Q) which is a measure of the separation of trait clusters which are clusters of traits that covary among themselves independently of others (He at al., 2020). We used the *igraph* package (Csardi & Nepusz, 2006) to extract network edge density (ED) and modularity (Q), while functional variability shape (FS) was calculated as the variance of the eigenvalues of the full trait-correlation matrix.

#### Phylogenetic history and/or environment effects on population-level coordination patterns

We characterised population-specific trait coordination by estimating, for each population, a Spearman trait–trait correlation matrix using individual-level data (Supplementary material, Fig. 7). Pairwise dissimilarity among populations was quantified using Manhattan distance, which penalizes both differences in magnitude and opposite correlation signs. The dissimilarity matrix was normalized to its maximum and converted into a similarity matrix (1 − dissimilarity). Population similarity patterns were visualized using hierarchical clustering. The effects of phylogenetic structure, aridity, and their interaction on population-level coordination patterns were tested using PERMANOVA directly on the dissimilarity matrix, with significance assessed using 9,999 permutations.

All statistical analyses were performed in R (version 4.5.2, R Core Team, 2025)

## Results

### Genetic clustering

Hierarchical clustering based on the distance matrix revealed a clear structure among populations. At k = 2, populations were separated into a West European group and a second group including Central and East European populations. At k = 3, FRA00005 emerged as a distinct cluster within the West European group. At k = 4, which was the best-supported clustering solution, Central and East European populations were further separated. However, to avoid single-population clusters and ensure robust statistical comparisons, FRA00005 was retained within the West European group in following analyses. We found significant differences in distance structure among the three groups (F = 4.21, R² = 0.34, p = 0.001), with group identity explaining 34.5% of the total variation. These three groups correspond to major geographic regions: West, Central, and East Europe (Fig. 2; Supplementary Material, Fig. 4).

### Magnitude of trait variability

We found the lowest among-population CV for P50 (5.41%), TLP (7.81%) and wood density (4.07%) (Fig. 3). Within populations wood density, slope and Ks CV was higher compared to among populations CV. In contrast, SMRI, g_res_ and HV exhibited the highest among-population CV (52.4%, 30.6 and 33.7%, respectively), accompanied by wide ranges of within-population CV (16.1–78.4% for g_res_, 22.2–68.8% for SMRI and 19.5–68.7% for HV).

**Figure 3.**
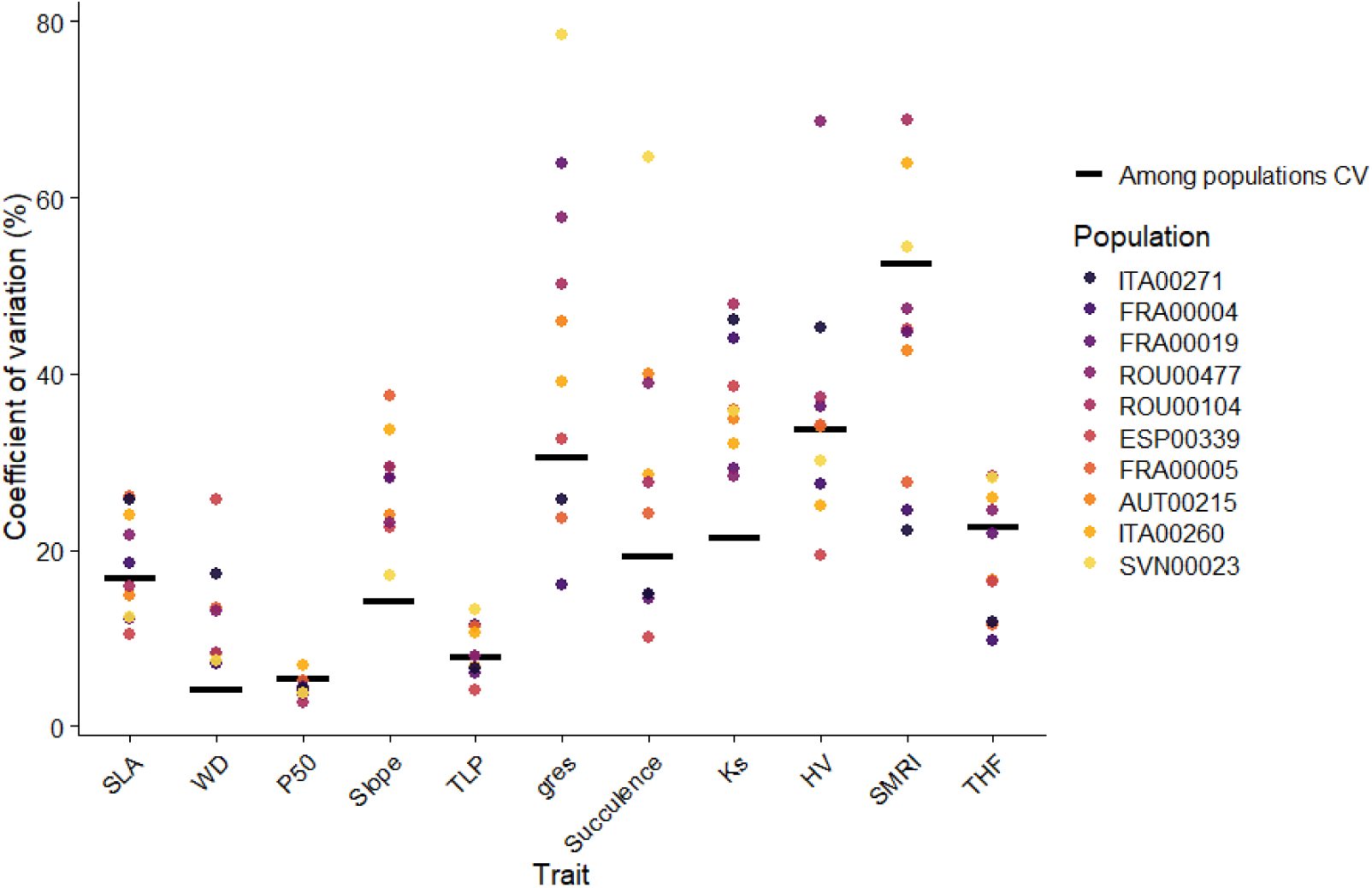
Coefficient of variation (CV) for each population (coloured dots) and among populations (solid black line) of *A. alba*. Colour points show each population which are sorted in decreasing aridity order (black to yellow dots). in the legend

### Phenotypic adjustment and environmental effect on within-population trait variation

The fixed predictors (age, aridity index or precipitation of the warmest quarter, and competition) explained on average 12 % of the variance of trait values across all full hierarchical models, with lowest variance for K_s_ and HV and highest for THF, SMRI and g_res_ (Fig. 4). Across traits, phylogenetic structure accounted on average between 2.7 and 74.6% of the variance depending on the trait, with the highest contribution for K_s_ and HV and moderate contribution to P50 and TLP (Fig. 4), while for slope, succulence and THF the credible interval of explained variance included zero (Table 4, Supplementary Material). Variance among populations within phylogenetic groups accounted on average for 1.9-37.8% with the highest contribution for SLA and THF, while for others traits the credible interval included zero (slope and succulence; Table 4, Supplementary Material). Residual (within-population and among population not phylogenetic) variance represented the largest component for most traits, particularly wood density, slope, g_res_ and succulence (Fig. 4).

**Figure 4.**
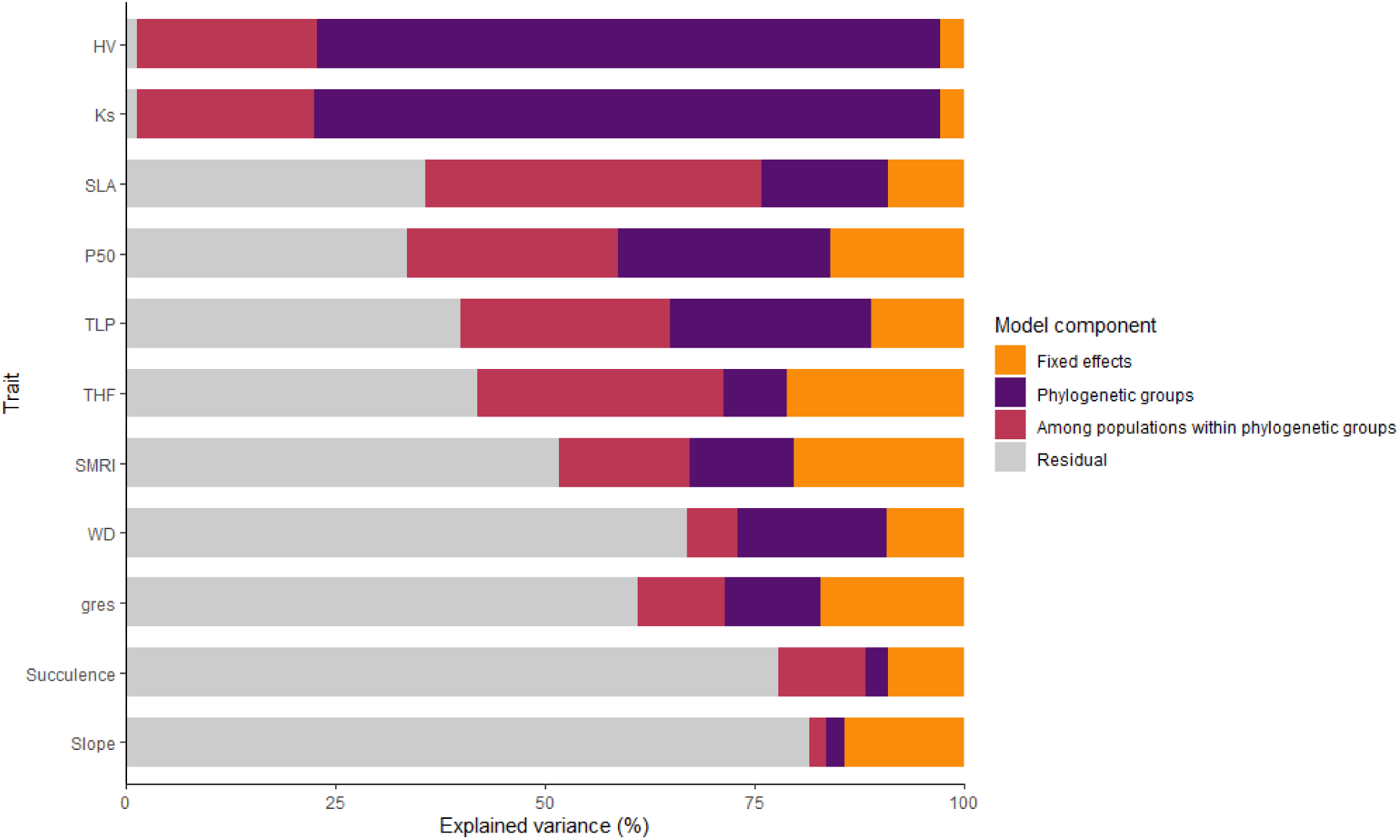
Partitioning of variance in functional trait values across full hierarchical Bayesian models. Bars show the mean percentage of variance explained by fixed effects (age, aridity index or precipitation of the warmest quarter, and tree competition), phylogenetic groups, variation among populations within phylogenetic groups, and residual variance (within-population and non-phylogenetic among-population variation). Confidence intervals are shown in supplementary material, table 4.

We found that tree age showed very strong evidence of a positive association with wood density and the slope of vulnerability curves (standardized estimate mean, estimate hereafter = 2.58, 2.65, respectively) and strong evidence of a positive association with SLA (estimate=1.82). Tree competition showed a moderate negative effect for P50 (estimate= −1.47), indicating more negative values of P50 in trees undergoing stronger competition). The aridity index exhibited moderate positive effects on the slope (estimate= 1.60), TLP (estimate= 1.26) and succulence (estimate= 1.25), indicating that populations from more humid environments (higher aridity index values) tended to show steeper vulnerability slopes, less negative values of TLP and higher succulence. Precipitation of warmest quarter showed a very strong effect for g_res_ (estimate= 2.47), SMRI (estimate= −2.27) and THF (estimate= −2.04) and a positive moderate effect for P50 (estimate= 1.18), indicating that trees in populations where precipitation of warmest quarter is higher showed higher g_res_, less negative values of P50 and lower SMRI and THF (Fig. 5).

**Figure 5.**
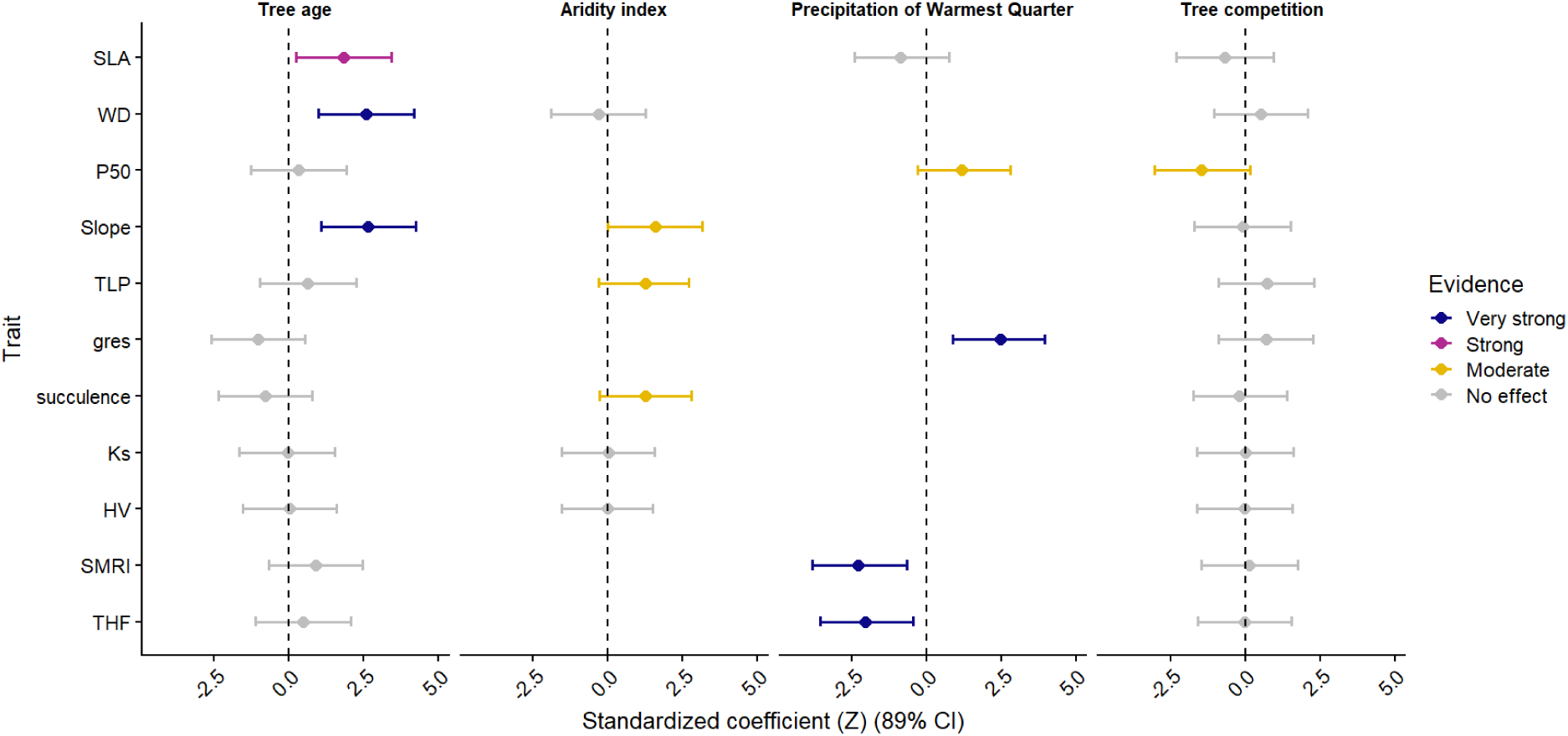
Standardised coefficient of Bayesian models (mean and 89% CI) examining the relationship of tree age, aridity index, precipitation of warmest quarter and tree competition, and each trait. Colour represents the strength of evidence for an effect according to the probability of direction (PD) with grey PD ≤ 0.9 = no evidence of effect; yellow 0.9 < PD < 0.95 = moderate evidence; pink 0.95 < PD < 0.975 = strong evidence; blue PD > 0.975 = very strong evidence

For traits without correlation between population mean and population SD (Supplementary material, Fig. 5), we found a positive significant relationship between aridity index and g^res^ SD and succulence SD, indicating greater variability in both traits in less arid sites (Fig. 6a, c). No significant correlations between SD or CV and the aridity index were found for the other traits (Fig. 6).

**Figure 6.**
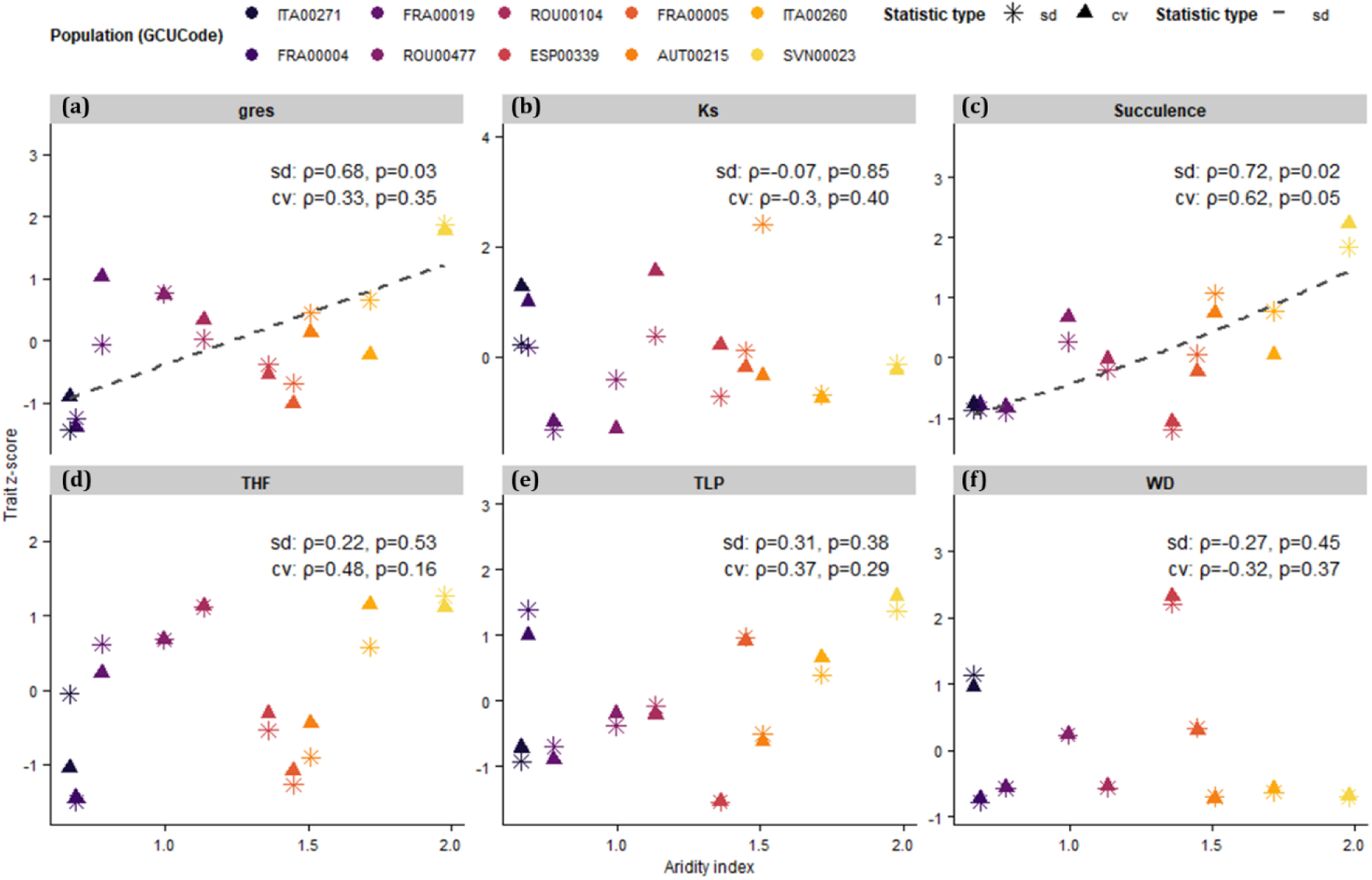
Rank correlations between trait standard deviation (SD) or trait coefficient of variation (CV) and aridity index for: g_res_ (a), Ks (b), succulence (c), THF (d), TLP (e) and WD (f). SD and CV values were z-scored to improve comparability among statistic types. Coloured points represent populations and different shapes correspond to statistic type (SD, or CV). Lines show fitted relationships obtained using generalized additive models (GAM), displayed only when Spearman’s rank correlations were significant (p < 0.05). Coefficient of correlation is shown (ρ)

### Trait coordination among and within populations

Within-population networks showed a low edge density (ED = 12.7%) and a low functional shape (FS = 0.75), indicating that individual-level trait combinations are only weakly coordinated and distributed across many small modules (7 modules, modularity = 0.16). In contrast, among-population networks were more structured, with a higher edge density (ED = 16.4%) and more than double functional shape (FS = 1.83). The among-population network also showed substantially higher modularity (Q = 0.65) and fewer modules (4) (Fig. 7). Among populations, we found a positive correlation between P50 and K_s_ and P50 and WD, and a negative correlation between TLP and HV. Within populations, we found a positive relationship between P50 and TLP and between g_res_ and leaf succulence.

**Figure 7.**
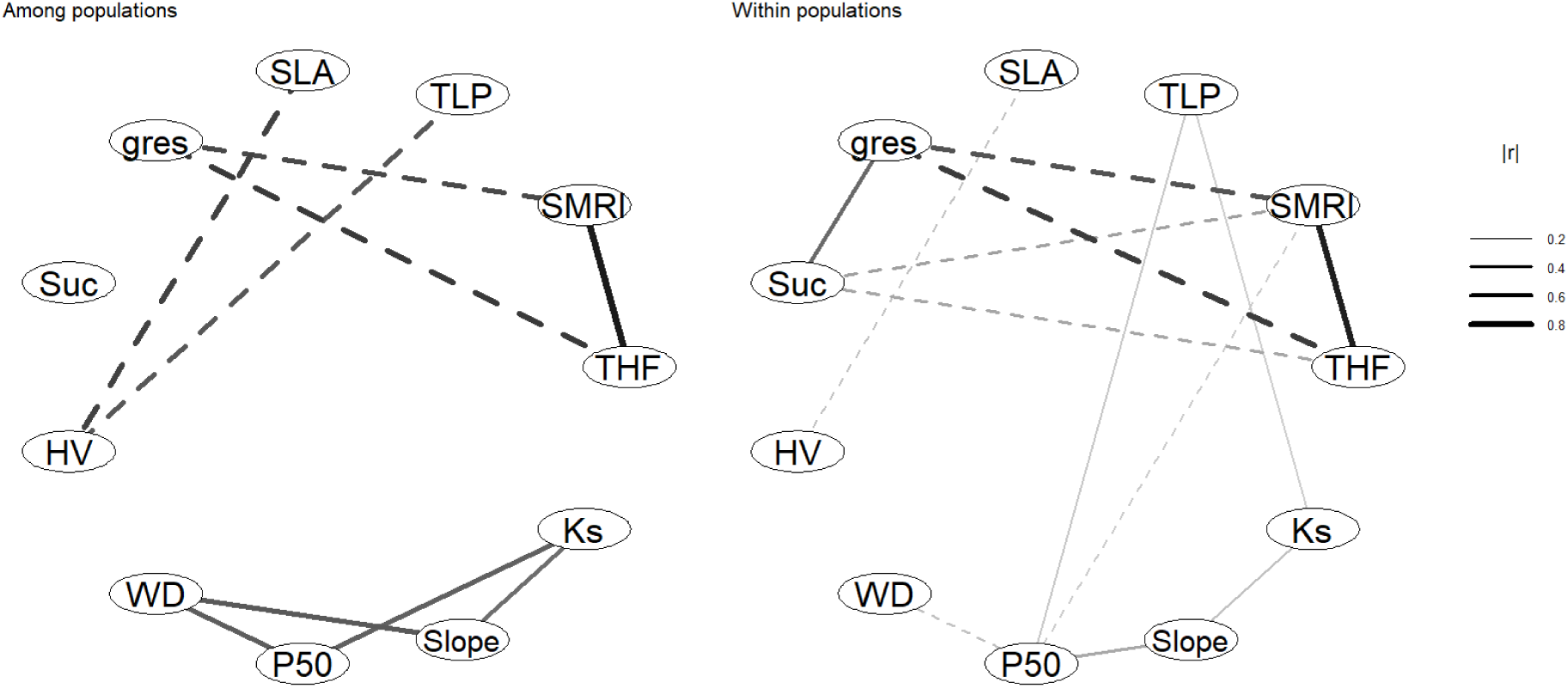
Trait correlation networks among (right panel) and within populations (left panel) of *A. alba* based on rank correlations. Solid lines represent significant positive correlations and dashed lines are significant negative correlations. Both line width and colour intensity scale with the absolute strength of the correlation.

### Phylogenetic history and/or environment effects on population-level coordination patterns

Network representations of hydraulic trait coordination are shown for each population (Fig. 7; Supplementary Material), providing a visual overview of trait associations within populations, organized by phylogenetic group and aridity index. Population-level hydraulic trait coordination was significantly structured by the interaction between phylogenetic structure and aridity (R² = 0.63, p = 0.037; Fig. 8, Supplementary Material). In contrast, neither phylogenetic structure nor aridity alone explained coordination patterns.

## Discussion

### Drivers of phenotypic adjustment and within-population phenotypic variance of hydraulic traits

By combining phylogenetic structure, large-scale climatic variables and within-population factors, our study provides an integrated assessment of how these sources of variation contribute to shaping phenotypic adjustment in hydraulic traits in silver fir. Populations in more arid sites exhibited lower slopes of vulnerability curves and indicated greater turgor maintenance (more negative values of TLP). Moreover, populations experiencing low precipitation during the warmest quarter showed reduced vulnerability to cavitation, residual leaf conductance and greater safety margin retention index and time to hydraulic failure. Interestingly, succulence, which is often interpreted as a proxy for internal water storage and thus associated with survival during drought (Blackman et al., 2019; Peters et al., 2021), was higher in humid populations of silver fir.

Drought resistance strategies can be separated into those that delay the onset of water stress (e.g. avoidance through stomatal regulation and water storage) and those that confer tolerance to severe water stress through hydraulic traits (Delzon 2015). Our results suggest that *A. alba* populations deploy different combinations of these strategies across environments. Trees from arid populations appear to rely primarily on drought tolerance, maintaining turgor and hydraulic function to lower water potentials (drought tolerance strategy) while minimising post-closure water losses through low residual leaf conductance (avoidance strategy). This combination likely allows trees to endure prolonged dry periods before hydraulic failure. In contrast, trees from wetter populations may mitigate drought stress through greater internal water storage, which can buffer short-term variability in water availability but may be less effective under sustained drought. It has to be noticed that our approach cannot fully disentangle trait responses to current environmental conditions from legacy effects of past environmental events, which may reflect acclimation processes.

For the two traits associated with the avoidance strategy (g_res_ and succulence), we found that higher local aridity reduced within-population phenotypic variance and coefficients of variation, a pattern similar to that reported by Fririon et al. (2023) for growth sensitivity to drought. This reduction in variance, combined with a shift towards lower trait values in more arid sites, indicates that arid conditions constrain the expression of hydraulic avoidance traits. Similar to the continental-scale patterns across species reported by Peters et al. (2021), in which aridity has acted as a strong selective pressure shaping hydraulic safety margins, our results suggest that increasing aridity restricts flexibility and imposes stronger constraints on these traits. This pattern supports the idea that marginal populations of *A. alba* may exhibit genetic signatures of adaptation to local climatic extremes, as also suggested by Roschanski et al. (2016), although in their study the strongest signal was associated with cold rather than aridity. However, it remains uncertain whether populations from wetter sites can adjust g_res_ values in response to increasing drought stress, either through phenotypic plasticity or evolutionary adaptation. A limited capacity to reduce residual water loss could increase the risk of hydraulic failure under future drought conditions in these currently wetter populations. It should be noted that restricting the analysis to traits lacking mean–variance relationships may be overly conservative, as such patterns can arise from the joint action of stabilising selection (modulating variance) and local adaptation (shifting trait means), as may be the case for traits exhibiting phenotypic adjustment. On the other hand, within-population variance in WD, TLP, K_s_ and THF was unrelated to their mean values, suggesting that differences in within-population variability among populations may be influenced by other factors, although these drivers were not captured by the climatic variables considered in this study (Fig. 3, supplementary material).

At the within-population scale, phenotypic adjustment was partly driven by tree age and competition. Age significantly influenced SLA, wood density and vulnerability-curve slope, whereas competitive environment affected P50. This highlights the importance of intrinsic factor and microenvironmental heterogeneity in modulating hydraulic function at the individual-tree level. The relevance of within-population sources of variation agrees with results found in *Pinus sylvestris*, where local competition and age influence key functional traits (Benavides et al., 2021). Meanwhile increasing neighbouring basal area has been suggested to buffer negative impact of drought events (Decarsin et al., 2024), we found that it increases xylem vulnerability to cavitation. This result could represent an opportunity to improve xylem resistance through forest management strategies such as reducing basal area at the local scale, although such interventions may also increase soil temperature and evaporative demand and alter selective pressures on regeneration. Therefore, these trade-offs should be considered when designing management strategies in *A. alba* populations.

We found a strong variance explained by phylogenetic signals for hydraulic efficiency traits such as HV and K_s_ which could indicate a high evolutionary conservatism in these traits (Sanchez-Martinez et al., 2020). Meanwhile, other traits related to drought response and hydraulic safety (TLP, P50, g_res_) showed high unexplained residual variation which could reflect phenotypic plasticity, microenvironmental heterogeneity and/or fine-scale genetic variation within-populations. Thus, our study supports that different evolutionary and ecological constraints affect each hydraulic trait.

### Magnitude of variation for hydraulic traits

We found that trait variability in *A. alba* spans a wide range, some traits being well conserved whereas others are more variable. Moreover, the variation of each trait is also different among populations resulting in higher variability within some populations compared to the species-level variability.

The lowest variability was found in P50, TLP, and wood density. At broader interspecific scales, hydraulic safety margins tend to converge, with most tree species operating close to their cavitation thresholds (Choat et al., 2012). Similarly, at the intraspecific level, P50 typically shows limited phenotypic variability and low plasticity (Martínez-Vilalta et al., 2009; Lamy et al., 2014), as observed in the present study, which may reflect underlying genetic or physiological constraints. The low variability observed in TLP has also been documented previously (Rosas et al., 2019). This limited variation may reflect functional constraints, since TLP represents a key physiological threshold in plant dehydration responses (Browne et al., 2023), and changes in this trait may increase mortality risk (Henry et al., 2019). The conservation of these traits suggests strong functional constraints in the capacity of *A. alba* to shift its hydraulic safety margin. We found also a low variability in wood density, which is not a direct hydraulic safety trait but has been shown to be related to drought safety (Rosner, 2017). Its low intraspecific variability in *A. alba* has been found previously. Fajardo et al. (2022), suggested the existence of phylogenetic and biomechanical constraints in wood density, which may indirectly restrict the range of hydraulic adjustments available to the species.

In contrast, *A. alba* showed high variability in g_res_. This is consistent with the relatively high plasticity of this trait, which has been reported to occur over very short timescales within individuals (Burlett et al., 2025), as well as over medium- and long-term periods (Duursma et al., 2019; Wang et al., 2025). We also found a relatively high variability in other traits, such as HV and Ks, or the drought resistance indices computed from multiple traits (SMRI and THF). Similar patterns have been reported for other species, where higher variability has been attributed to the integrative nature of some traits, as they involve multiple physiological processes (Siefert et al., 2015; Rosas et al., 2019).

### Coordination of traits is strong among populations

Scale dependence in trait coordination has important implications for understanding drought responses in *A. alba*. Our network analyses revealed a clear contrast between scales: trait coordination was weak and fragmented within populations, whereas it was stronger and more modular among populations. A similar but reversed scale dependence has been reported for other functional traits in trees (Alonso & Herrera, 2001), and together these results suggest that the structure of trait covariation depends strongly on both scale and trait type, which could reflect different underlying biological processes. At the among-population level, coordinated trait networks likely represent the outcome of past evolutionary processes, integrating long-term selection under contrasting environmental conditions and population genetic differentiation, consistent with the influence of large-scale climate and phylogenetic structure (Ohsawa & Ide, 2008). In contrast, within-population correlations describe phenotypic covariation on which selection may act in the future, representing potential constraints for adaptive responses to drought (Bolnick et al., 2011; Westerband et al., 2021). Thus, our results suggest that at the within-population level, *A. alba* is less constrained for future adaptive responses to drought than suggested by the among-population trait coordination networks.

This scale dependence was particularly evident for hydraulic safety–efficiency relationships. The safety–efficiency trade-off can be evaluated through the relationships between hydraulic efficiency traits (K_s_ or HV) and hydraulic safety thresholds (P50 or TLP) (Fuchs et al., 2021). Among populations, less negative P50 was associated with higher K_s_, and populations with less negative TLP exhibited lower HV values, consistent with increased hydraulic efficiency at the expense of safety (Jacobsen et al., 2007; Gleason et al., 2016; Markesteijn et al., 2011). Although these trade-offs have often been attributed to climatic differences among populations (Rosas et al., 2019), we found no clear association with site aridity or past drought intensity (Fig. 7 and 8; Supplementary Material), suggesting that population genetic differentiation in hydraulic efficiency traits contributes to shaping these patterns. In contrast, safety–efficiency trade-offs were absent within populations. Instead, P50 and TLP were positively related, indicating coordinated stomatal regulation at the individual level, whereby stomatal closure is adjusted to embolism resistance to maintain hydraulic safety margins (Martin-StPaul et al., 2017). Similarly, the positive within-population correlation between residual leaf conductance and leaf succulence suggests short-term compensatory strategies at the individual level. The absence of these relationships among populations reinforces the interpretation that within-population correlations reflect functional adjustments rather than outcomes of past selection.

Overall, trait coordination among populations reflects the legacy of past evolution, whereas within-population trait networks describe the constraints and capacities for future adaptation, underscoring the importance of explicitly accounting for scale when interpreting functional trait relationships. This multilevel perspective is consistent with broader integrative frameworks that emphasise combining physiological, genetic and ecological dimensions to assess species’ vulnerability to climate change (Williams et al., 2008).

## Conclusions

Our study demonstrates that intraspecific variation in hydraulic and structural traits of *Abies alba* reflects the combined effects of environmental conditions, phylogenetic structure and within-population processes acting at different scales. Hydraulic safety traits (P50, turgor loss point and wood density) were highly constrained across the species range, indicating limited capacity to adjust hydraulic safety margins under increasing drought. In contrast, traits related to water-use regulation and hydraulic efficiency were more variable, revealing greater potential for phenotypic adjustment. Environmental gradients and phylogenetic structure primarily shaped trait differentiation and coordinated safety–efficiency trade-offs among populations, reflecting the legacy of past selection. Within-populations trait variation and coordination were mainly driven by intrinsic factors such as tree age, competition, defining the phenotypic covariation on which future selection may act. Together, these results highlight the importance of scale in interpreting trait variability and coordination, and suggest that although *A. alba* may partly adjust to increasing drought through flexible traits, strong constraints on key hydraulic safety traits could limit adaptive responses under future climate change.

## Acknowledgements

This research was funded by the French ANR-22-CE02-0010-01 project FLORES and by European Union (EU)’s Horizon 2020 Research and Innovation Program N. 862221 “FORGENIUS”. The authors wish to thank Lucas Antunes, Camilla Avanzi, Didier Besombes, Marie-Claude Bouhedi, Marc Busuldu, Gaëlle Capdeville, Marianne Corréard, Jonathan Feichter, Vega Garcia-Segura, Damien Gounelle, Sonia Hernando, Arnaud Jouineau, Nicolas Mariotte, Marion Parizat, Andrea Piotti, Flaviu Popescu, Dragos Postolache, Joan Prunera-Olivé, Adolfo Sanmartín-Arévalo, Daniel Suciu and Léa Veuillen as well to the whole FORGENIUS consortium for support in collecting samples

## Authors contributions

B.A-M, F.L. and N.M.-S.P. designed the research. M.M., F.J., H.C., S.D., J.M.T-R. and N.M.-S.P. conducted the field and/or lab work. I.S. analysed genetic data and A.D. carried out sensibility analyses. B.A.M analysed general results with inputs from F.L., N.M.-S.P., A.C., B.F. and C.S.-S. B.A- M. produced a first draft of the manuscript with inputs from F.L, N.M.-S.P. and A.C. All authors reviewed the manuscript.

## Data availability

All data will be made publicly available in an appropriate repository upon acceptance of the manuscript, and the corresponding link will be provided.

