## Supplementary Material for "Multiscale intraspecific variation and coordination of hydraulic traits in silver fir"

**Table 1**. Population code, Lat and Log, (Latitude and Longitude, respectively, decimal degrees), Elevation (m), Slope (degree), BA_Intra, BA_Inter (Basal Area Intra- and Interspecific, respectively, m2/ha), AI (aridity index) and other species present in the stand (Other spp)

| Code | Lat | Long | Elevation | Slope | BA_Intra | BA_Inter | AI | Other spp |
|---|---|---|---|---|---|---|---|---|
| ITA00271 | 38.14 | 15.86 | 1718 | 18 | 13.21 | 37.54 | 0.6642 | Fagus sylvatica |
| FRA00004 | 41.98 | 9.11 | 1573 | 25 | 64.97 | 0 | 0.6878 | |
| FRA00019 | 44.18 | 5.24 | 1147 | 34 | 15.51 | 12.55 | 0.7777 | Acer opalus, Fagus sylvatica, Sorbus aria |
| ROU00477 | 47.64 | 24.01 | 924 | 30 | 18.06 | 27.55 | 0.9951 | Acer platanoides, Fagus sylvatica, Picea abies |
| ROU00104 | 45.26 | 24.08 | 1366 | 19 | 13.15 | 15.57 | 1.1335 | Fagus sylvatica |
| ESP00339 | 42.69 | -0.11 | 1350 | 14 | 18.60 | 43.84 | 1.3602 | Fagus sylvatica, Ilex aquifolium, Pinus sylvestris, Populus tremula, Taxus baccata |
| FRA00005 | 46.85 | 5.99 | 770 | 0 | 18.58 | 6.74 | 1.4471 | Crataegus monogyna, Picea abies |
| AUT00215 | 47.45 | 14.70 | 1180 | 11 | 16.05 | 32.10 | 1.5086 | Fagus sylvatica, Picea abies |
| ITA00260 | 44.36 | 10.09 | 1350 | 37 | 23.10 | 29.82 | 1.7149 | Fagus sylvatica, Picea abies, Sorbus aucuparia |
| SVN00023 | 45.62 | 14.47 | 776 | 14 | 21.78 | 12.98 | 1.9774 | Fagus sylvatica, Picea abies, Ulmus |

**Table 2**. Mean ± SD of 11 study traits in each population

| GCUCode | gres | Succulence | TLP | P50 | Slope | Ks | HV | SLA | WD | SMRI | THF |
|---|---|---|---|---|---|---|---|---|---|---|---|
| **AUT00215** | 2.48 ± 1.14 | 329.02 ± 131.70 | 2.29 ± 0.16 | 3.20 ± 0.13 | 167.29 ± 40.25 | 6.17e-04 ± 2.15e-04 | 1.72e-04 ± 5.83e-05 | 5.31e-03 ± 7.87e-04 | 0.55 ± 0.04 | 0.43 ± 0.18 | 72.13 ± 12.71 |
| **ESP00339** | 2.25 ± 0.73 | 238.30 ± 24.05 | 2.36 ± 0.10 | 3.86 ± 0.15 | 108.31 ± 24.47 | 3.37e-04 ± 1.30e-04 | 1.93e-04 ± 3.76e-05 | 5.30e-03 ± 5.58e-04 | 0.47 ± 0.12 | 0.74 ± 0.33 | 82.13 ± 15.34 |
| **FRA00004** | 1.94 ± 0.31 | 273.16 ± 41.08 | 2.27 ± 0.26 | 3.48 ± 0.14 | 113.57 ± 31.95 | 3.51e-04 ± 1.55e-04 | 9.35e-05 ± 2.58e-05 | 7.79e-03 ± 1.45e-03 | 0.52 ± 0.04 | 0.67 ± 0.16 | 84.86 ± 8.36 |
| **FRA00005** | 2.49 ± 0.59 | 347.45 ± 83.79 | 2.11 ± 0.24 | 3.63 ± 0.19 | 130.86 ± 49.00 | 4.26e-04 ± 1.53e-04 | 1.00e-04 ± 3.42e-05 | 8.54e-03 ± 2.22e-03 | 0.51 ± 0.07 | 0.66 ± 0.18 | 79.38 ± 10.03 |
| **FRA00019** | 1.40 ± 0.89 | 266.49 ± 38.78 | 2.42 ± 0.15 | 3.55 ± 0.16 | 148.59 ± 41.91 | 3.90e-04 ± 1.14e-04 | 2.25e-04 ± 8.19e-05 | 5.31e-03 ± 6.50e-04 | 0.53 ± 0.04 | 1.06 ± 0.48 | 103.13 ± 23.70 |
| **ITA00260** | 3.18 ± 1.25 | 412.53 ± 117.99 | 1.96 ± 0.21 | 3.66 ± 0.26 | 155.77 ± 52.44 | 4.08e-04 ± 1.31e-04 | 1.44e-04 ± 3.59e-05 | 6.21e-03 ± 1.49e-03 | 0.52 ± 0.04 | 0.68 ± 0.44 | 77.43 ± 23.44 |
| **ITA00271** | 0.84 ± 0.22 | 266.41 ± 40.13 | 2.02 ± 0.13 | 3.69 ± 0.16 | 122.91 ± 36.20 | 3.39e-04 ± 1.56e-04 | 9.94e-05 ± 4.50e-05 | 7.09e-03 ± 1.82e-03 | 0.52 ± 0.09 | 2.11 ± 0.47 | 145.88 ± 18.86 |
| **ROU00104** | 1.87 ± 0.94 | 258.52 ± 71.45 | 2.24 ± 0.18 | 3.39 ± 0.10 | 133.00 ± 39.11 | 3.35e-04 ± 1.60e-04 | 1.60e-04 ± 5.99e-05 | 6.49e-03 ± 1.04e-03 | 0.52 ± 0.04 | 0.78 ± 0.54 | 90.63 ± 27.31 |
| **ROU00477** | 2.25 ± 1.30 | 241.23 ± 93.72 | 2.02 ± 0.16 | 3.34 ± 0.12 | 121.32 ± 28.00 | 4.88e-04 ± 1.39e-04 | 1.17e-04 ± 8.04e-05 | 6.29e-03 ± 1.37e-03 | 0.50 ± 0.07 | 0.78 ± 0.37 | 90.88 ± 24.15 |
| **SVN00023** | 2.35 ± 1.84 | 260.99 ± 168.70 | 1.97 ± 0.26 | 3.48 ± 0.13 | 141.85 ± 24.29 | 4.10e-04 ± 1.46e-04 | 9.05e-05 ± 2.73e-05 | 6.80e-03 ± 8.49e-04 | 0.53 ± 0.04 | 0.92 ± 0.50 | 94.88 ± 28.43 |

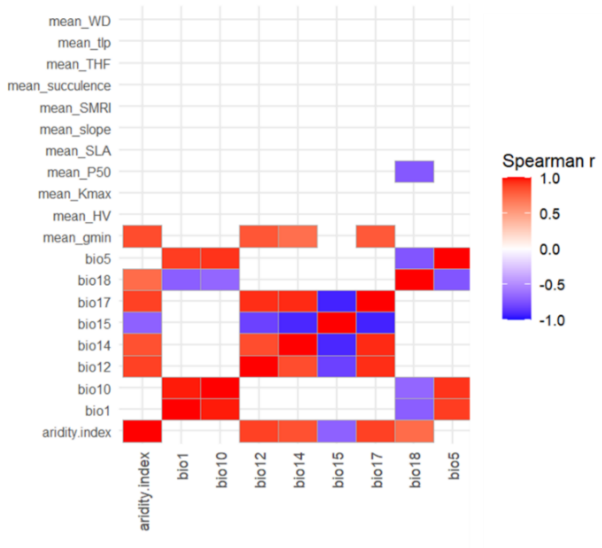


**Figure 1.** Rank correlations between mean trait for each population and different climatic variables. Bio1: Annual Mean Temperature; bio5 = Maximum Temperature of Warmest Month; bio10 = Mean Temperature of Warmest Quarter; bio12 = Annual Precipitation; bio14 = Precipitation of Driest Month; bio15 = Precipitation Seasonality (Coefficient of Variation); bio17 = Precipitation of Driest Quarter; bio18 = Precipitation of Warmest Quarter

**Table 3.** Percentage of variance (mean and credible interval) explained by fixed in two alternative Bayesian models: one with competition, tree age and aridity index (model A) and other with competition, tree age and precipitation of the warmest quarter (model B).

| Trait | Model A | Model B |
|---|---|---|
| SLA | 9.1 (1.2–22.5) | 10.1 (1.5–24.6) |
| WD | 9.1 (1.9–18.6) | 8.8 (1.8–17.9) |
| P50 | 10.4 (0.9–25.3) | 15.7 (1.7–34.7) |
| slope | 14.2 (4.9–24.6) | 12.9 (4.2–23.5) |
| TLP | 11.2 (1.0–27.8) | 6.3 (0.5–17.7) |
| gres | 16.9 (2.6–34.3) | 17.1 (3.2–32.6) |
| succulence | 8.7 (1.3–20.3) | 6.2 (0.8–15.2) |
| Ks | 2.7 (0.0–11.5) | 2.2 (0.0–9.4) |
| HV | 2.7 (0.0–11.3) | 2.2 (0.0–8.9) |
| SMRI | 12.7 (1.5–30.3) | 20.2 (3.6–37.9) |
| THF | 16.4 (1.7–37.2) | 21.1 (2.9–42.1) |

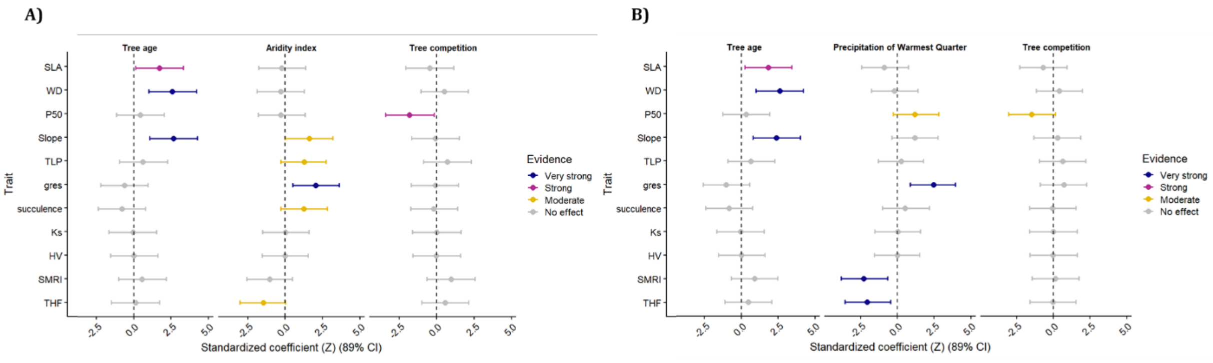


**Figure 2**. Standardised coefficient of Bayesian models (mean and 89% CI) of two alternative models for each trait. Panel (A) shows the effects of tree age, aridity index and tree competition and panel (B) shows the effects of tree

age, precipitation of the warmest quarter and tree competition each trait. Colour represent the strength of evidence for an effect according to the probability of direction (PD) with grey PD ≤ 0.9 = no evidence of effect; yellow 0.9 < PD < 0.95 = moderate evidence; pink 0.95 < PD < 0.975 = strong evidence; blue PD > 0.975 = very strong evidence.

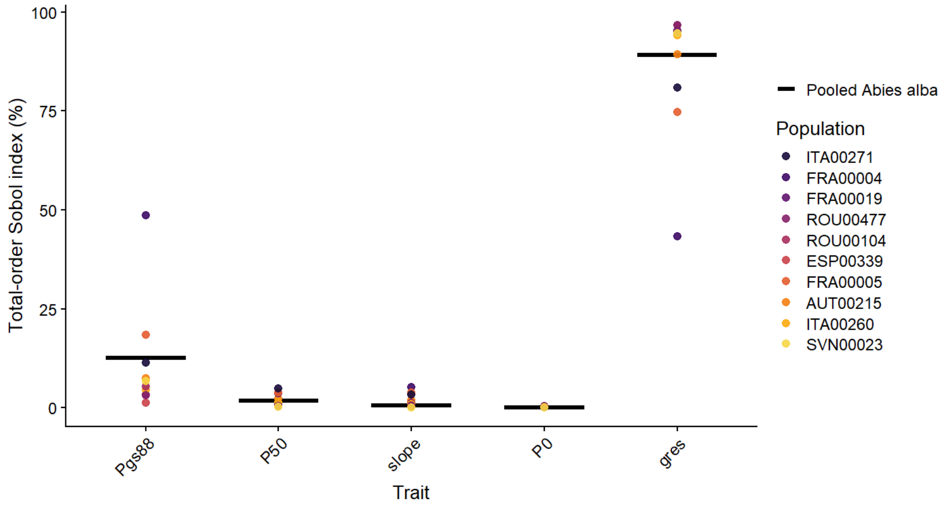


**Figure 3.** Relative importance of hydraulic traits in determining variation in the SurEau model output (THF) across *A. alba* populations based on total-order Sobol sensitivity indices. Points represent the relative contribution (%) of each trait to THF variance for each population, while the black horizontal line indicates the value obtained for the pooled *A. alba* dataset. Water potential causing 88% stomatal closure (Pgs88); Osmotic potential at full turgor of the leaf symplast (P0).

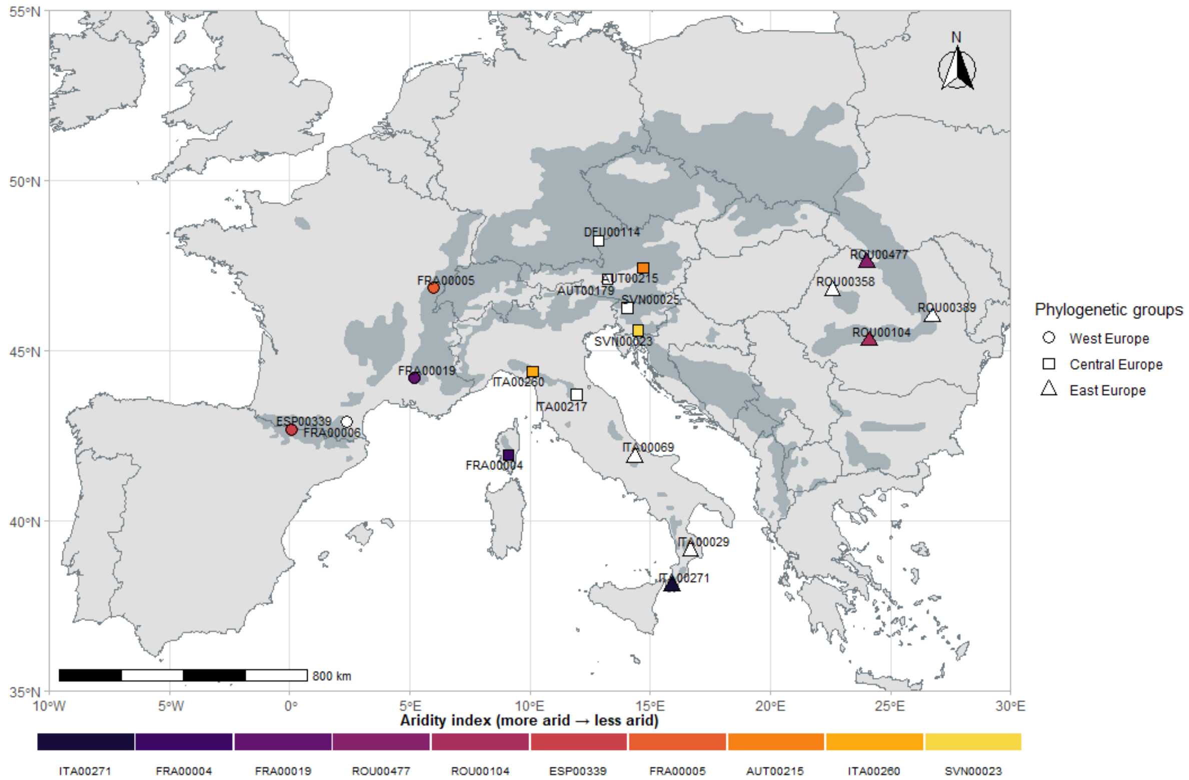

**Figure 4.** Geographic distribution of *A. alba* populations across Europe. Coloured symbols represent the aridity index for the 10 populations selected for trait measurements, whereas non-coloured symbols correspond to additional populations included for genetic analyses. Symbol shapes denote phylogenetic groups (West Europe, Central Europe, and East Europe). The natural distribution range of *A. alba* is shown in dark grey.

**Table 4.** Mean and credible interval of variance attributable to the proportion of total variance accounted by the fixed predictors, phylogenetic cluster groups component (phylogenetic groups), among populations within phylogenetic groups population component (among populations within phylogeographic groups) and within-population variance (residual).

| Trait | Fixed effects | Phylogenetic groups | Among populations within phylogenetic groups | Residual |
|---|---|---|---|---|
| SLA | 10.1 (1.4–23.9) | 14.7 (1.0–56.3) | 37.8 (15.1–62.8) | 37.3 (13.5–59.9) |
| WD | 9.2 (2.0–18.6) | 18.0 (2.0–61.1) | 5.9 (1.1–15.9) | 66.8 (31.1–85.9) |
| P50 | 15.6 (1.6–34.4) | 26.3 (1.7–78.2) | 24.5 (3.9–53.1) | 33.5 (9.4–55.0) |
| Slope | 14.1 (4.9–24.7) | 2.7 (0.0–12.5) | 1.9 (0.0–10.1) | 81.3 (65.5–93.1) |
| TLP | 11.1 (1.0–27.2) | 24.1 (1.4–76.3) | 24.7 (4.0–51.8) | 40.1 (11.1–65.3) |
| gres | 17.3 (3.3–32.7) | 11.5 (0.3–50.2) | 10.5 (0.5–31.4) | 60.7 (31.4–81.1) |
| Succulence | 8.9 (1.3–20.8) | 2.8 (0.0–15.4) | 9.9 (0.0–33.0) | 78.4 (49.1–96.2) |
| Kmax | 2.7 (0.0–10.9) | 75.2 (39.7–98.0) | 20.8 (1.6–51.5) | 1.3 (0.1–3.2) |
| HV | 2.7 (0.0–10.8) | 74.8 (39.4–98.0) | 21.2 (1.7–52.4) | 1.3 (0.1–3.3) |
| SMRI | 20.4 (3.4–38.4) | 12.4 (0.3–54.8) | 15.7 (1.1–38.9) | 51.5 (23.9–73.3) |
| THF | 21.2 (2.9–42.3) | 6.5 (0.0–37.0) | 29.9 (9.4–56.8) | 42.3 (20.3–63.0) |

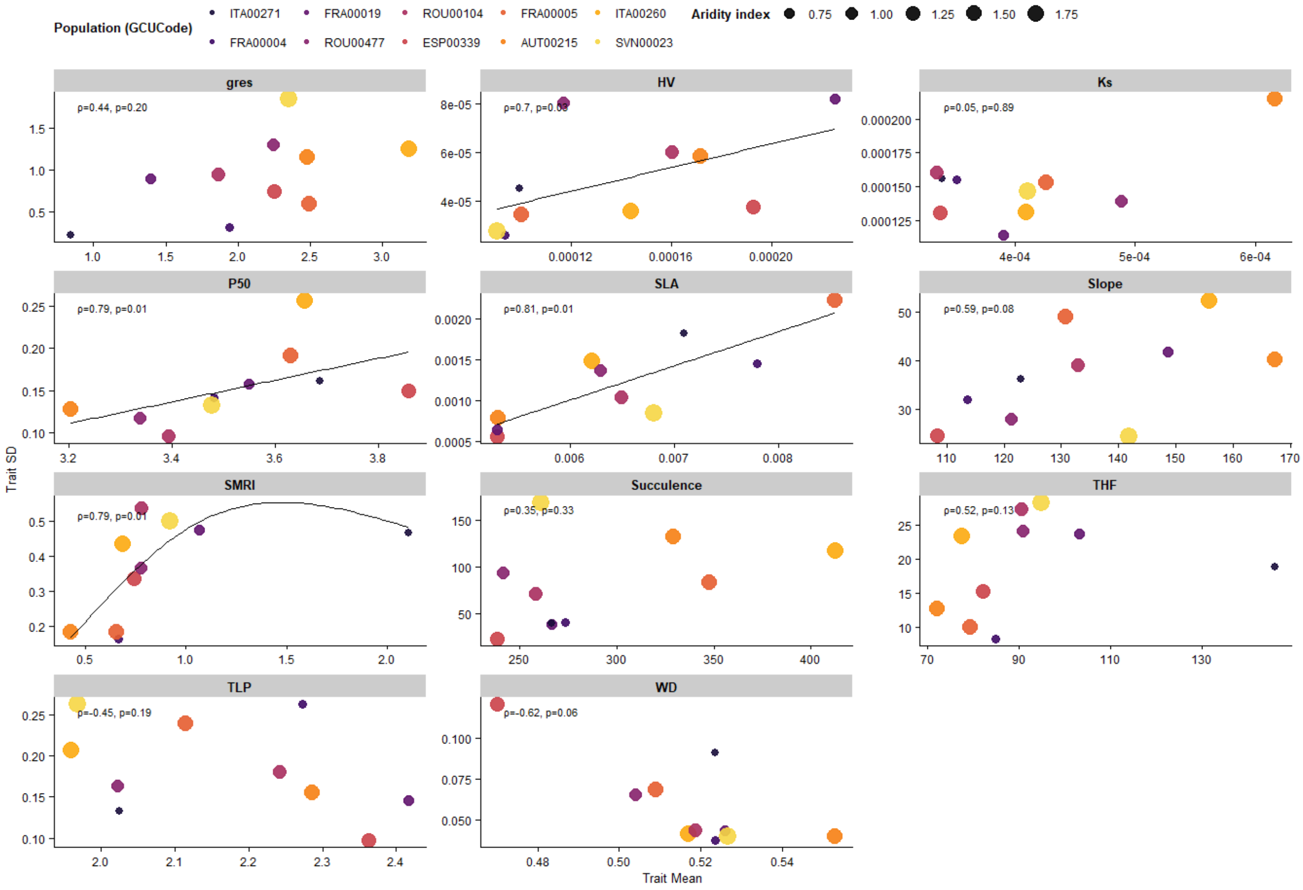


**Figure 5.** Rank correlations between trait mean and trait SD for all traits. Point size reflects the aridity index (larger symbols indicate less arid populations). Lines show fitted relationships obtained using generalized additive models (GAM), displayed only when Spearman's rank correlations were significant (p < 0.05).

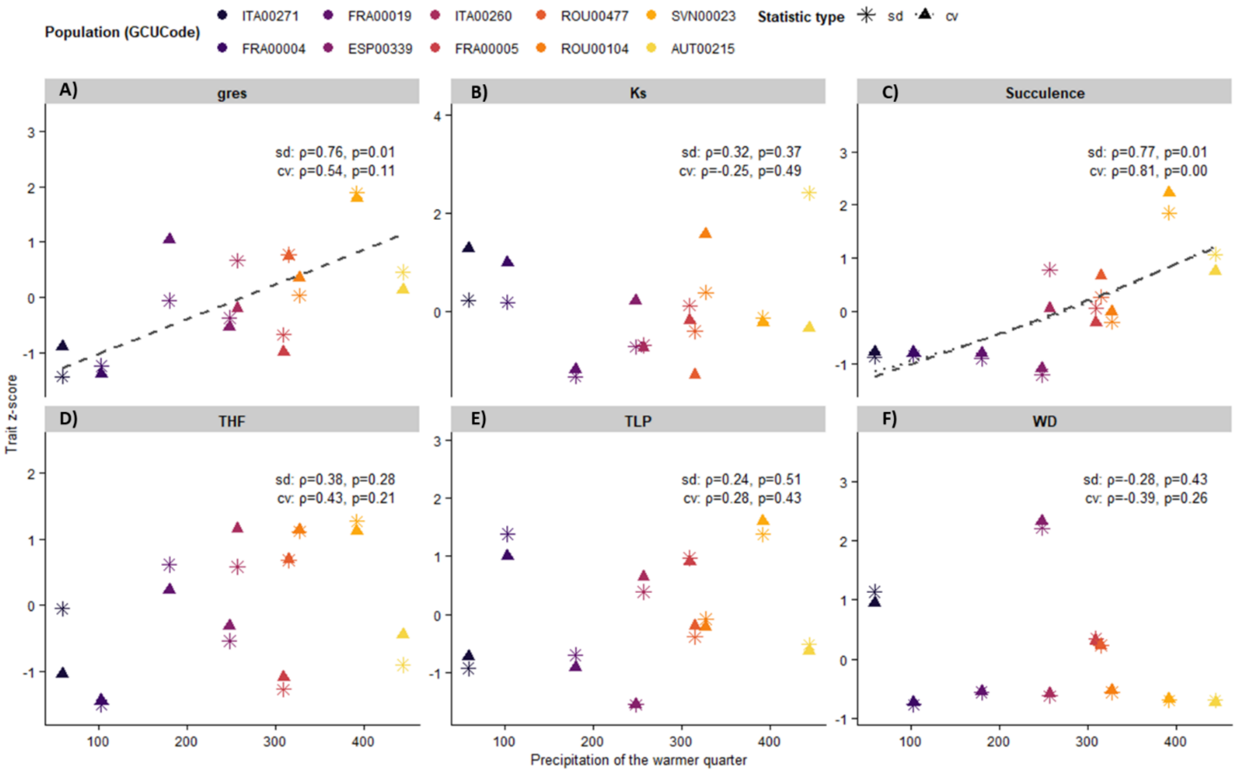


**Figure 6**. Rank correlations between trait SD or trait CV and precipitation of the warmest quarter for: gres (A), Ks (B), succulence (C), THF (D), TLP (E) and WD (F). SD and CV values were z-scored to improve comparability among statistic types. Coloured points represent populations and different shapes correspond to statistic type (SD, or CV). Lines show fitted relationships obtained using generalized additive models (GAM), displayed only when Spearman's rank correlations were significant ($p < 0.05$).

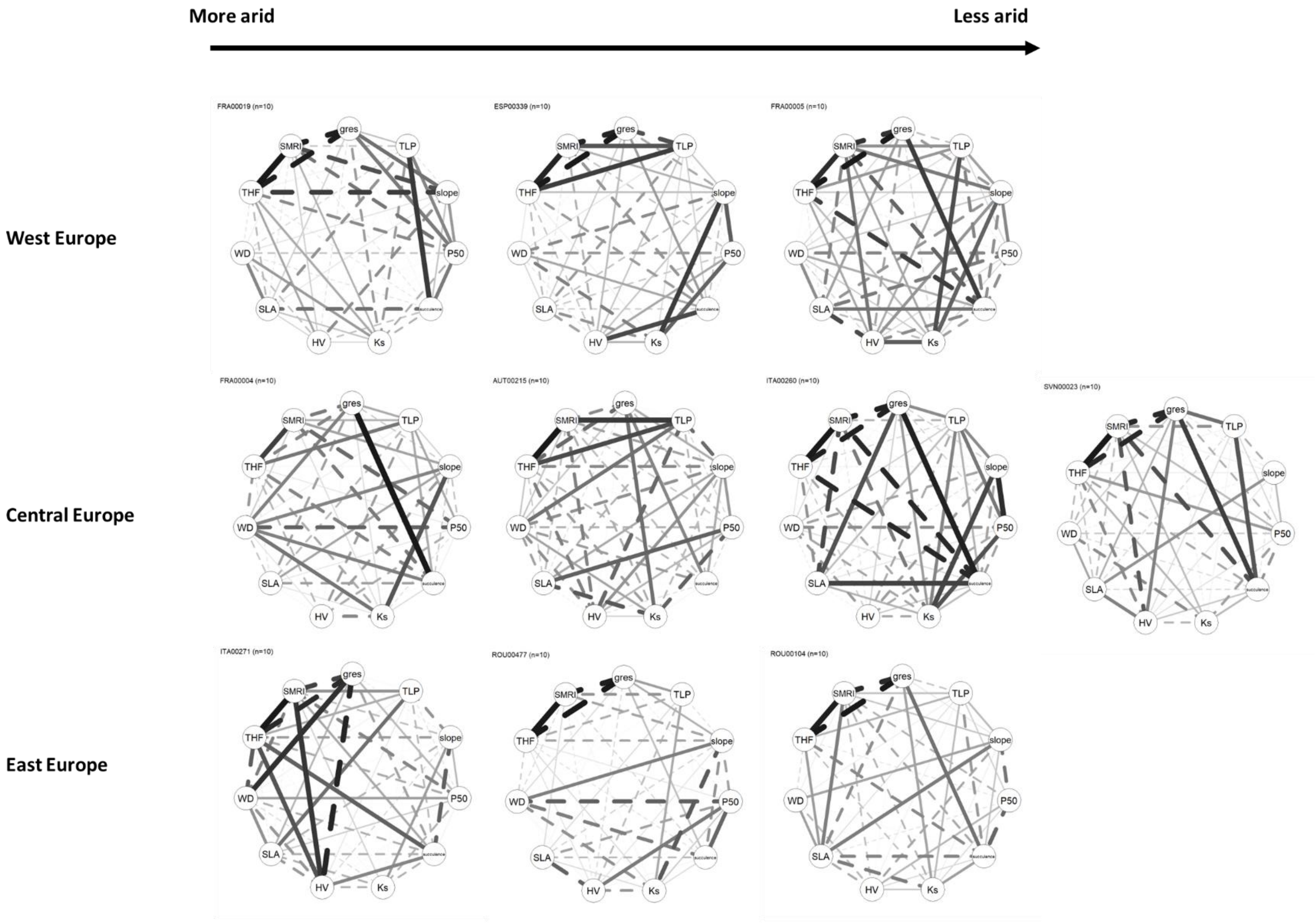


**Figure 7.** Trait correlation networks among individuals within each population of *A. alba* based on rank correlations. Populations are ordered based on aridity index (AI) for each phylogenetic group. Solid lines represent positive correlations and dashed lines are negative correlations. Both line width and colour intensity scale with the absolute strength of the correlation.

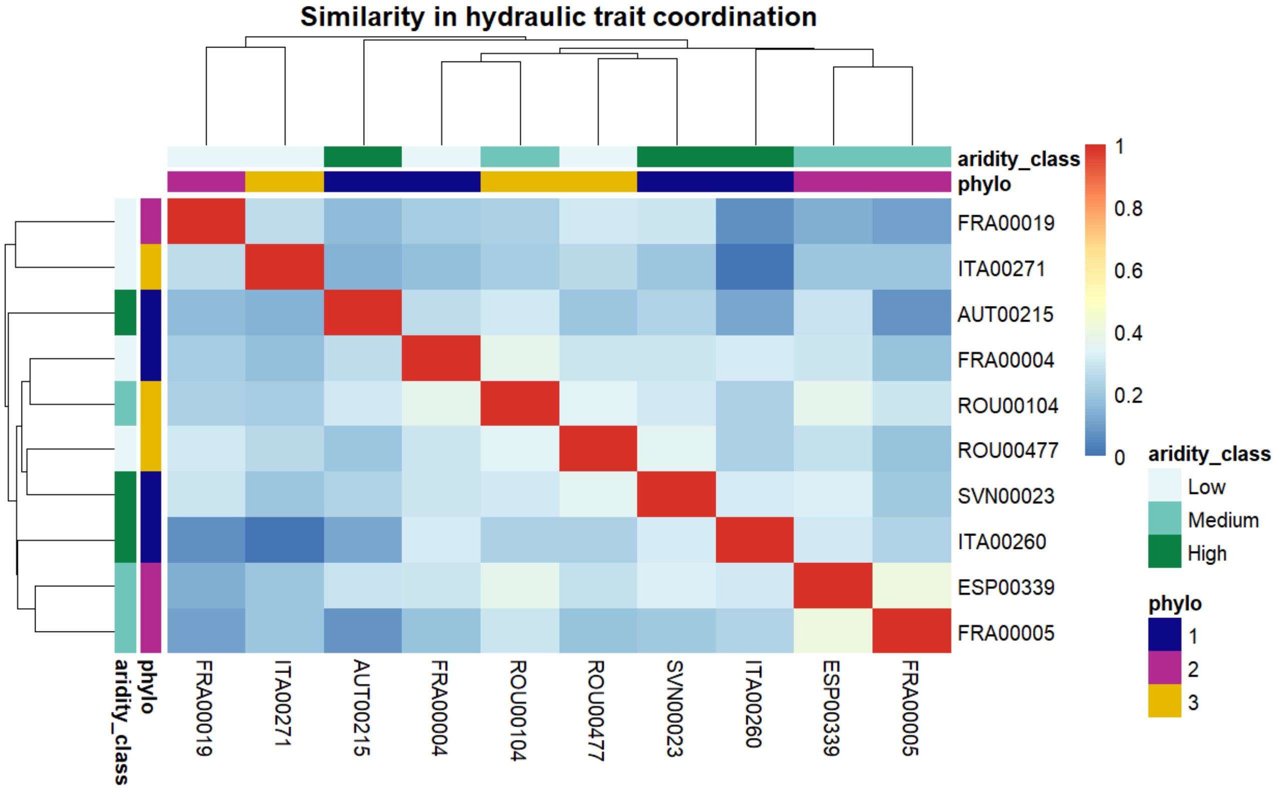


**Figure 8.** Heatmap of pairwise disimilarity in functional trait syndromes among populations, where light colours represent greater similarity. Hierarchical clustering of populations is based on the similarity matrix, and colour bars indicate aridity class (aridity index is divided in three class for a better representation) and phylogenetic groups.

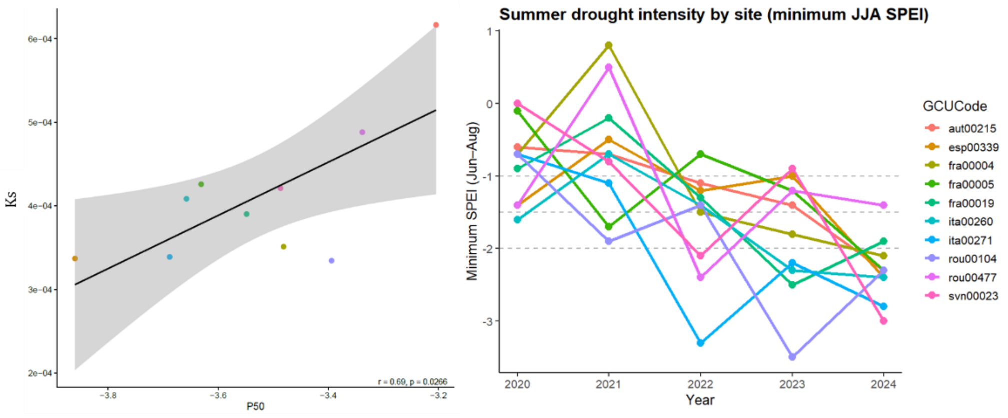


**Figure 9.** Correlation between P50 and Ks (left panel) and minimum SPEI of each population (right panel).

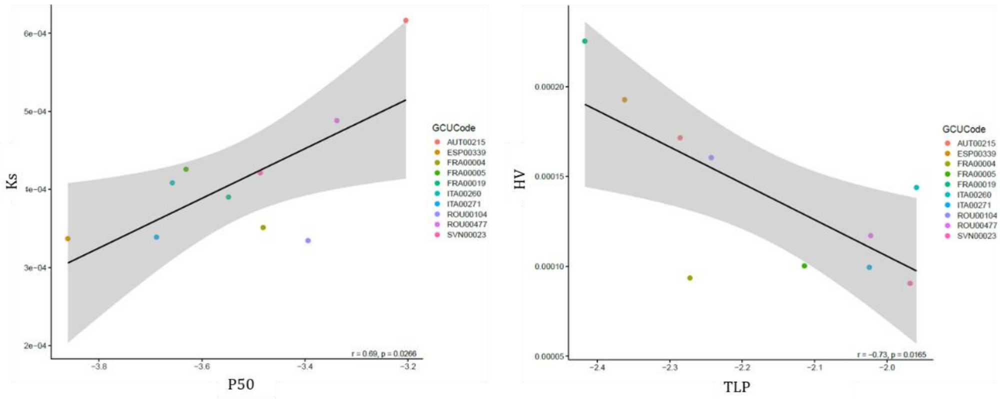


**Figure 10**. Correlation of population values between Ks and P50 (left panel) and HV and TLP (right panel).
